# Deciphering Cell Cycle Dynamics and Cell States in Single-cell RNA-seq data with SPAE

**DOI:** 10.64898/2026.03.05.709782

**Authors:** Jiahao Yi, Jiajia Liu, Peng Guo, Yuan-Nong Ye, Xiaobo Zhou

## Abstract

Rapid advances in single-cell RNA sequencing (scRNA-seq) technology have enabled the investigation of gene expression changes at the single-cell level, particularly for elucidating the heterogeneity among cells and complex biological processes. This technique reveals subtle molecular differences within individual cells, thereby offering a unique viewpoint for the investigation of cell cycle progression, cellular differentiation, and disease pathogenesis. However, accurately identifying and analyzing cell cycle dynamics in scRNA-seq data remains challenging due to the complexity of the data and the subtle differences between cell states. To address this challenge, we developed the integrated Sinusoidal and Piecewise AutoEncoder (SPAE), an autoencoder-based piecewise linear model, for characterizing the cell cycle dynamics and cell states in scRNA-seq data. Compared with existing methods, SPAE demonstrates substantially improved accuracy and robustness in cell cycle characterization. Additionally, SPAE can accurately predict cancer cell cycle transitions and effectively facilitate the removal of cell cycle effects from gene expression data. SPAE is available for non-commercial use at https://github.com/YaJahn/SPAE.

## Introduction

In recent years, the field of biomedical research has experienced significant advancements due to the development of single-cell RNA sequencing (scRNA-seq) technology [1–3]. This technology represents a significant transition from tissue level to single-cell level, providing a unique perspective for a deeper understanding of cell heterogeneity [4]. Moreover, scRNA-seq has demonstrated immense potential in studying two fundamental biological processes: the cell cycle and cell differentiation [5–7].

The cell cycle, a fundamental framework in cellular fate, is essential for organismal growth and development. Its stages, G1, S, G2, and M phases, are interconnected by complex molecular events and regulatory networks. Precise regulation of these stages is crucial for maintaining tissue homeostasis and enabling developmental adaptation [8, 9]. Moreover, it is intricately associated with cell states and plays a pivotal role in tumorigenesis. Consequently, accurate identification and comprehension of cell cycle stages are imperative for the in-depth exploration of cellular behaviors [10, 11].

scRNA-seq enables precise quantification of gene expression at the individual-cell level, thereby offering novel and comprehensive insights into the cell cycle [12, 13]. Despite the rich datasets provided by scRNA-seq for cell cycle analysis, the interpretation of these data presents several challenges. A primary hurdle is the accurate inference of specific cell cycle stages from the scRNA-seq data. Traditional experimental methods, though able to identify cell cycle stages, are not only time-consuming and labor-intensive but also lack the capability for the quantitative measurement of cell cycle phase [14]. Moreover, the technical variability and data sparsity inherent to scRNA-seq [15], compounded by the transient and overlapping nature of cell cycle stages, significantly complicate the analysis [16]. This complexity underscores the need for advanced methodologies to accurately interpret cell cycle dynamics from scRNA-seq data.

Addressing the challenges associated with scRNA-seq data interpretation, the scientific community has recently introduced several computational methodologies. These encompass supervised machine learning methods, which predominantly utilize known cell cycle genes. For instance, cyclone [5] and the CellCycleScoring function in Seurat [17] are notable examples. These methods employ annotated cell cycle genes for predicting the cell cycle phases (G1, S, or G2/M) of individual cells. Additionally, reCAT [18] is an innovative approach combining the Travelling Salesman Problem and Hidden Markov Model to reconstruct cell cycle pseudotime series. However, a significant limitation of these methods is their dependency on datasets with pre-annotated cell cycle genes and experimental labels, which constrains their broader application.

To circumvent these constraints, unsupervised techniques have been developed. CCPE [19] is a representative linear autoencoder-based model for cell cycle analysis. It projects single-cell gene expression profiles into a low-dimensional latent space using a linear encoder and reconstructs the input with a linear decoder. A helical structure is then fitted in the latent space to capture the continuous cyclic trajectory of the cell cycle. While CCPE performs well when the data distribution is approximately linear, its linear encoder limits its ability to model nonlinear or multi-stage cell cycle transitions, which motivated the development of SPAE. Other unsupervised methods include Cyclum [20] and CYCLOPS [21]. Cyclum utilizes an autoencoder model that integrates both linear and non-linear elements within the hidden layer, aimed at inferring the pseudotime of cellular cyclical processes. Conversely, CYCLOPS employs a linear-projection autoencoder, mapping data onto a closed elliptical curve in a low-dimensional space, offering a fresh perspective in understanding cell cycle dynamics. However, it’s important to note that CYCLOPS, while designed to simulate circadian rhythms, incorporates complex operations such as square roots and division in its neural network, potentially posing challenges in optimization. Moreover, tools like cyclone and reCAT, while useful, do not effectively eliminate cell cycle effects from expression data, highlighting the need for continued advancements in this field.

In this study, we present a novel computational framework, the Integrated Sinusoidal and Piecewise AutoEncoder (SPAE), designed to concurrently analyze cell cycle dynamics and cell states from single-cell RNA-seq data. The motivation behind SPAE arises from the need to more accurately capture both the cyclic and piecewise linear characteristics of cellular processes observed in scRNA-seq data. Existing models such as CCPE [19] use a purely linear encoder, which limits their ability to represent complex nonlinear gene expression trajectories, while models like Cyclum rely on sinusoidal transformations, they cannot explicitly distinguish multiple cell states that deviate from a single smooth cycle. To overcome these limitations, SPAE integrates two complementary components: a nonlinear component to represent the periodic nature of the cell cycle and a piecewise linear component to model transitions between distinct cellular states that often follow locally linear patterns in gene expression space. This combination allows SPAE to more faithfully reconstruct cyclic and branching trajectories while simultaneously assigning cells to specific states. We rigorously assessed SPAE’s efficacy in estimating cell cycle pseudotime and determining cell stage classifications. Our comparative analysis includes established methodologies such as CCPE, cyclone, Seurat, Cyclum, CYCLOPS, and reCAT. Furthermore, we demonstrate SPAE’s utility in predicting cancer cell cycle transitions and in mitigating the confounding effects of cell cycle variations.

## Method

### Datasets

We utilized datasets presented in **Table 1** and **Table 2** to assess the performance of SPAE. The Quartz-Seq dataset of mouse embryonic stem cells (mESCs) [22], sequenced through Quartz-Seq technology, provided cell cycle stage and gene expression data for 33,412 genes. The H1 human embryonic stem cells (hESCs) [23] dataset utilized a fluorescence ubiquitination-based cell cycle indicator to stage 247 cells. For the E-MTAB-2805 mESCs dataset [6], 288 mouse embryonic stem cells were sequenced using the HiSeq 2000 sequencing system, covering 38,293 genes. To assess model robustness under different dropout rates, we utilized the E-MTAB-2805 mESCs dataset, which initially had a dropout rate of 24%. We first applied the MAGIC [24] model to impute the missing data, and subsequently introduced artificial dropout rates of 0%, 20%, 50%, and 70%. Additionally, the nutlin-treated multiple cancer cell lines dataset [25] included single-cell RNA-seq data from 24 cancer cell lines, treated with DMSO or nutlin on the 10× Genomics platform, highlighting cell cycle arrest induced by nutlin in cells expressing wild-type *TP53*. The FELINE Breast Cancer Single-Cell Genomics dataset [26] comprised patients with ER+ breast cancer undergoing neoadjuvant endocrine therapy (letrozole) with or without a CDK4/6 inhibitor (ribociclib), sampled at the start of treatment, after 14 days, and after 180 days of treatment, using 10× technology for single-nucleus RNA sequencing.

**Table 1.**
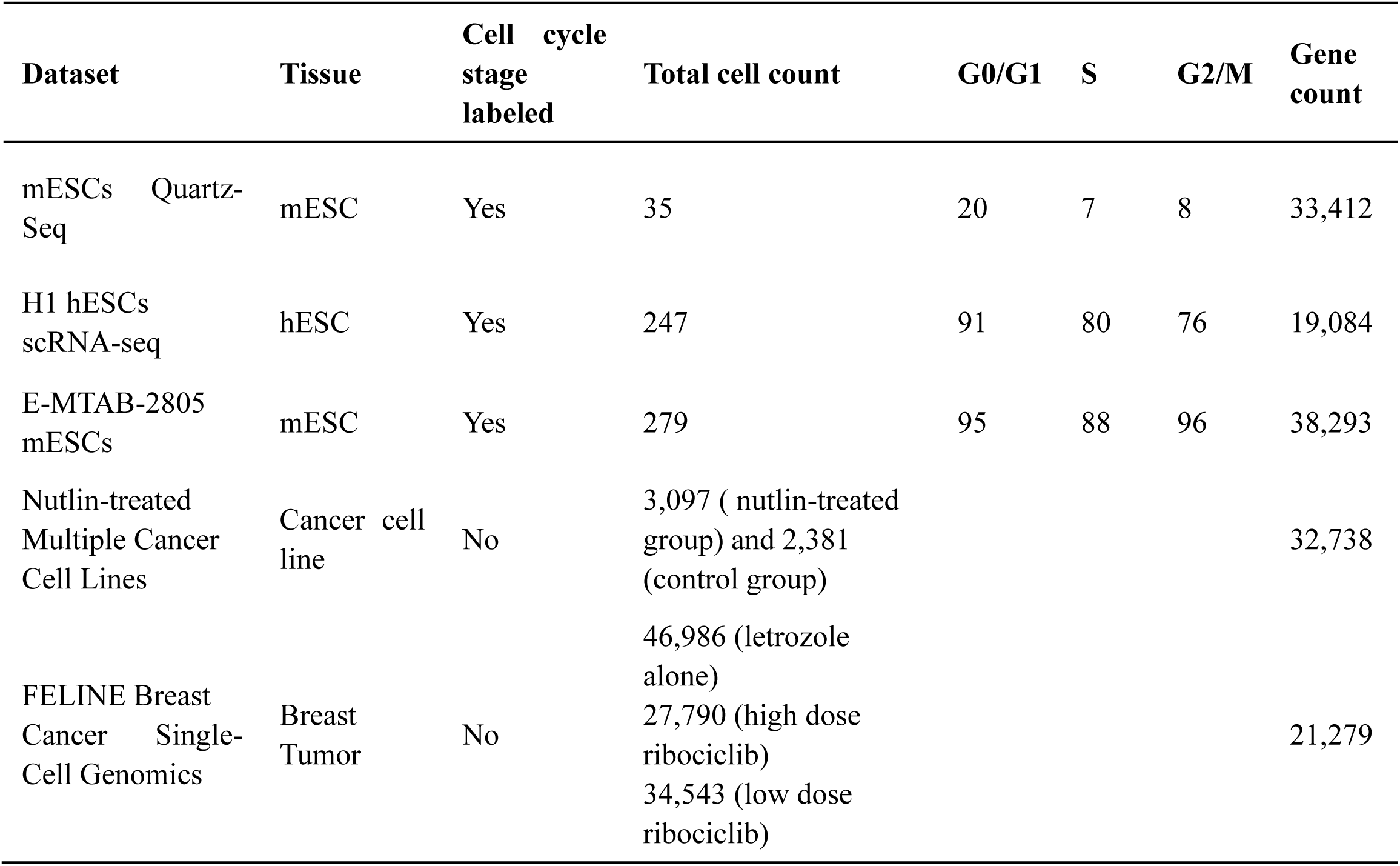
Cell-cycle scRNA-seq datasets.

**Table 2.** scRNA-seq datasets of cell cycle effect removal.

| **Dataset** | **Tissue** | **Cell type** | **Total cell count** | **Gene count** |
| --- | --- | --- | --- | --- |
| Mouse ES cells | Mouse embryonic stem cellss | 4 | 2717 | 24,046 |
| hMyo | Differentiating myoblasts (0 h, 24 h, 48 h, 72 h) | 4 | 372 | 47,192 |
| Breast cancer | Human Breast Cancer (MDA-MB-231) | 4 | 376 | 16,383 |

Moreover, to evaluate SPAE’s performance in removing cell cycle effects, we analyzed datasets from mouse embryonic stem cell, human myoblasts (hMyo), and breast cancer. The mouse embryonic stem cell dataset [27] utilized droplet microfluidics for high-throughput barcoding and RNA sequencing of individual cells. This dataset focused on the impact of leukemia inhibitory factor withdrawal. It included data from different stages of mouse embryonic stem cells at different stages: undifferentiated, and 2, 4, and 7 days post LIF withdrawal. The scRNA-seq data of human myoblasts [7], developed by Trapnell et al. (2014), included differentiating myoblasts sampled at various time points, namely at 0, 24, 48, and 72 hours. At 0 hours (0h), myoblast cells are actively proliferating and remain undifferentiated, with transcriptional profiles dominated by cell cycle and proliferation-associated genes. By 24 hours (24h), after switching to a differentiation-induction medium, the cells begin exiting the cell cycle and initiating early differentiation programs, marked by downregulation of cell cycle genes and upregulation of early myogenic markers like MYOD1. At 48 hours (48h), the cells show active differentiation with increased expression of muscle-specific genes such as MYOG (myogenin), reflecting a transition from proliferative myoblasts to early muscle precursors. By 72 hours (72h), the differentiation process is largely complete, with transcriptional profiles dominated by late myogenic markers like MYH genes, indicating mature muscle cells with a heterogeneous mix of late differentiation states. These time points correspond to critical stages in an experimental setup where the growth medium was changed to an induction medium, triggering the transition of proliferating myoblasts into a differentiation program. This process promotes the differentiation of myoblasts into more specialized cell types, allowing the capture of dynamic transcriptional changes associated with cell fate transitions. Breast cancer dataset [28] involves high throughput sequencing of MDA-MB-231 breast tumor cells, exploring the role of CSL (CBF1/RBP-Jkappa/Suppressor of Hairless/LAG-1) in cancer. It includes four cell types: CSLKO1 and CSLKO2 (CSL gene knockouts) and WT1 and WT2 (wild-type controls).

### SPAE model

SPAE models the distinct cell states using a piecewise linear regression framework. Piecewise linear regression is a modeling approach in which the relationship between variables is represented by multiple linear segments with different slopes across distinct regimes [29]. The mathematical formulation, including the continuity constraints and objective function, is detailed in **Supplementary Note 1**. In SPAE, we employ an autoencoder-based piecewise linear model in which the encoder consists of both nonlinear and piecewise linear components. In the nonlinear component, we use a standard multi-layer perceptron with hyperbolic tangent activation functions to map the transcriptome profile *X* to *z^c^* which represents the pseudotime along the cell cycle process (**Figure 1**). The piecewise linear component assigned cells into different clusters. Suppose we have *k* clusters, the gate function in our piecewise linear model determined which cluster a cell *x_n_* belongs to, defined as

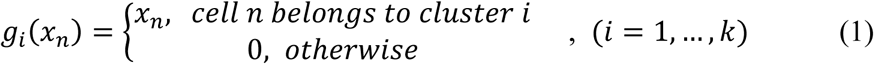

**Figure 1.**
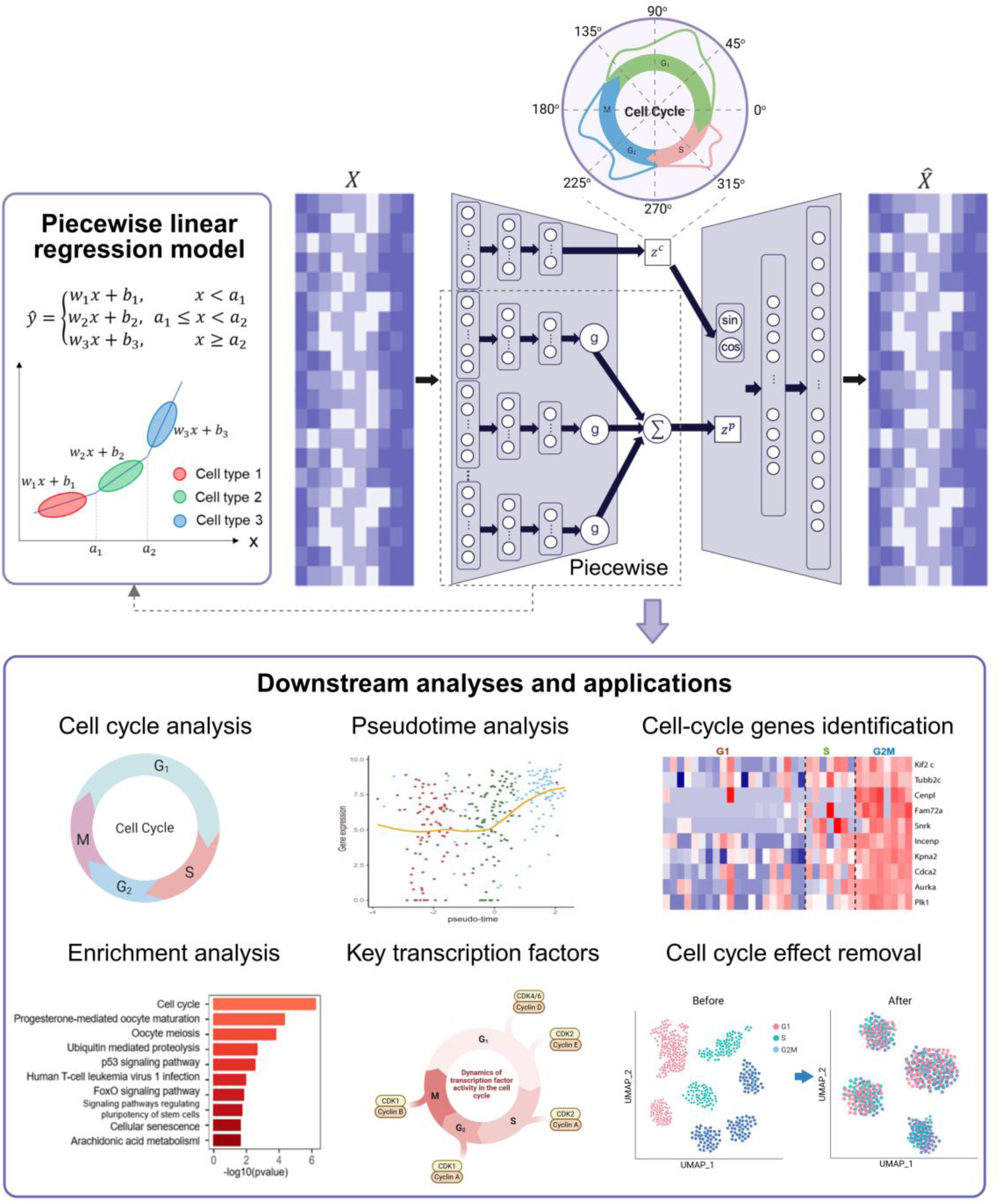
Overview of the SPAE framework. SPAE primarily consists of two components: a nonlinear component and a piecewise linear component. The nonlinear component is employed for the estimation of cell cycle pseudotime, while the piecewise linear component is dedicated to predicting various cell types. Six downstream analyses and applications of SPAE have been employed to evaluate its performance.

The transformations of the encoder can be represented as

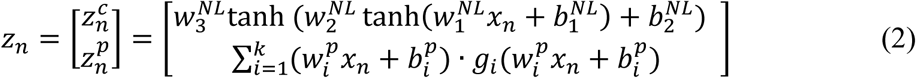

Where *w*^*NL*^ (collection of *w*_1_^*NL*^, *w*_2_^*NL*^, *w*_3_^*NL*^) and *b*^*NL*^ are the weight and bias matrices of the nonline encoder, *w*^*p*^ represents weight in the piecewise linear component. In the decoder, we used *V* as the weight matrix of the decoder and performed linear transformations, as follows

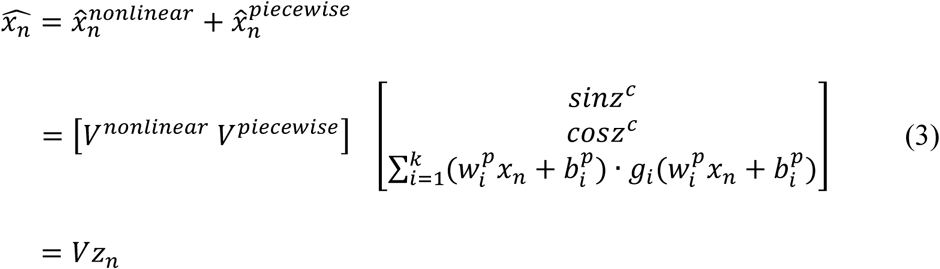

The optimization problem using the least square error is formulated as

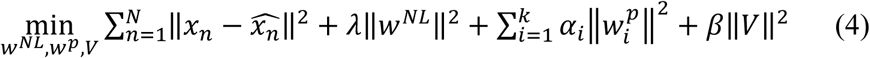

Where *λ*, *α*_*i*_, and *β* are regularization coefficients controlling the complexity of the nonlinear encoder, the piecewise linear component, and the decoder, respectively. Specifically, *λ* ∥*w^NL^*∥^2^ ensures the smoothness of the learned latent manifold by regularizing the nonlinear weights, while *α*_*i*_ constrains the slope parameters of each cell state. To train the model, we employ an alternating optimization strategy that iterates between refining the piecewise thresholds and updating the autoencoder weights. Detailed descriptions of the initialization and the two-step optimization algorithm are provided in **Supplementary Note 2**.

## Results

### Overview of SPAE

As illustrated in **Figure 1**, SPAE integrates an autoencoder to analyze single-cell RNA sequencing (scRNA-seq) data by capturing both cell cycle dynamics and different cell types. The model consists of two key components: a nonlinear encoder for cell cycle estimation and a piecewise linear regression model for identifying distinct cell types based on their gene expression profiles. In the encoder, a multi-layer perceptron with hyperbolic tangent activation functions reduces the data ’ s dimensionality while mapping cells along a pseudotime trajectory, capturing their progression through the cell cycle. To account for the periodicity of the cell cycle, SPAE employs sine and cosine functions in the decoder, allowing for precise estimation of pseudotime and cell cycle phases. Once the cyclic behavior is modeled, SPAE integrates a piecewise linear regression model, which allows the program to treat inferred cycle processes as confounding factors and, after discounting confounding cell cycle effects, to make predictions for multiple cell types.

### SPAE accurately infers cell cycle pseudotime

To assess the performance of SPAE in predicting cell cycle pseudotime, we compared SPAE with Cyclum, CYCLOPS, and reCAT [18, 20, 21], and the widely used trajectory inference tool Monocle using the scRNA-seq data of mESCs. **Figure 2A** shows the distribution of cell cycle pseudotime estimated by each method. We calculated statistical measurements using the interquartile distances between boxplots, which helped quantify the separation of pseudotime of cells in different cell cycle phases. Specifically, we measured the distance between the lower quartile of the inferred cell cycle pseudotime in the S phase and the upper quartile of the pseudotime in the G1 phase, as well as the distance between the lower quartile of the pseudotime in the G2/M phase and the upper quartile of the pseudotime in the S phase. Both SPAE and Cyclum retain the correct cell cycle order, from G1 to S and then to G2/M. CYCLOPS effectively distinguishes between S and G2/M phases, while reCAT can distinguish G1 and S phases but not G2/M. Compared to Cyclum, SPAE exhibits superior performance in separating S phase and G2/M phase. To quantitatively validate these observations, we calculated Spearman’s rank correlation coefficient (ρ)[30] between each method’s inferred pseudotime and the true, biologically ordered cell-cycle stages. Our results demonstrate that SPAE achieves the strongest monotonic correlation with the true cell-cycle order (ρ = 0.866, *P* = 1.90×10⁻¹¹), substantially outperforming other methods such as Cyclum (ρ = 0.699, *P*=2.96×10⁻⁶), reCAT (ρ = 0.591, *P* = 1.87×10⁻⁴), Monocle (ρ = 0.468, *P*=0.004587) and CYCLOPS (ρ = -0.276, *P* = 0.1087). Notably, while Monocle is effective for branching trajectories, its lower correlation here suggests limitations in capturing the closed-loop topology of the cell cycle compared to SPAE. These statistics confirm that SPAE most accurately reconstructs the biological sequence of cell-cycle stages, effectively capturing the continuous transition consistent with the visual patterns in Figure 2A. We calculated the Pearson correlation between the gene expression and cell cycle pseudotime inferred by SPAE. The genes with the highest correlation with cell cycle pseudotime were Aurora kinase A (*Aurka*), cell division cycle associated 2 (*Cdca2*) and karyopherin alpha 2 (*Kpna2*). The correlation coefficients of *Aurka*, *Cdca2*, and *Kpna2* are 0.73, 0.73, and 0.71, respectively (**Figure 2B**). *Aurka* is a kinase that plays an important role in cell cycle regulation and control. It affects the cell cycle mainly by regulating chromosome separation in the preparatory phase of cell division. *Aurka* maintains the stability of the cell cycle by interacting with other cell cycle proteins [31]. *Cdca2* has a crucial role in controlling the G1/S transition, which is a critical stage in the cell cycle. *Cdca2* depletion led to cell cycle arrest at the G1 phase, suggesting that *Cdca2* is required for proper cell cycle progression[32]. *Kpna2* is also involved in cell cycle regulation and restriction of cell cycle progression [33]. To complete the comparison, we extend this analysis to CYCLOPS, Cyclum, reCAT and Monocle, showing the Pearson correlation between gene expression and inferred cell cycle pseudotime for the top six genes identified by each method (**Supplementary Figure S1-S4**). Notably, most of the top genes identified by CYCLOPS, Cyclum, and reCAT have little to no direct involvement in the cell cycle, which suggests a potential limitation in these methods for capturing key cell cycle-related dynamics. Additionally, we included specific plots comparing the Pearson correlation for *Aurka*, *Cdca2*, and *Kpna2* between all methods. While SPAE consistently identified these genes as highly correlated with cell cycle pseudotime (with correlations of 0.73, 0.73, and 0.71, respectively), their correlations were significantly lower in CYCLOPS, Cyclum, and reCAT (**Supplementary Figure S1B-S3B**). For instance, Monocle showed weaker correlations for key regulators like *Aurka* (R = 0.55) and *Cdca2* (R = 0.58), although it maintained a comparable correlation for *Kpna2* (R = 0.77) (**Supplementary Figure S4B)**. This indicates that general-purpose tools like Monocle and other specific methods may fail to capture the importance of these well-known cell cycle regulators, further highlighting SPAE’s strength in identifying biologically relevant genes associated with cell cycle progression. **Figure 2C** displays a heatmap illustrating several G2/M phase marker genes with relatively high correlations to cell cycle pseudotime estimated by SPAE, which are all highly expressed in the G2/M phase.

**Figure 2.**
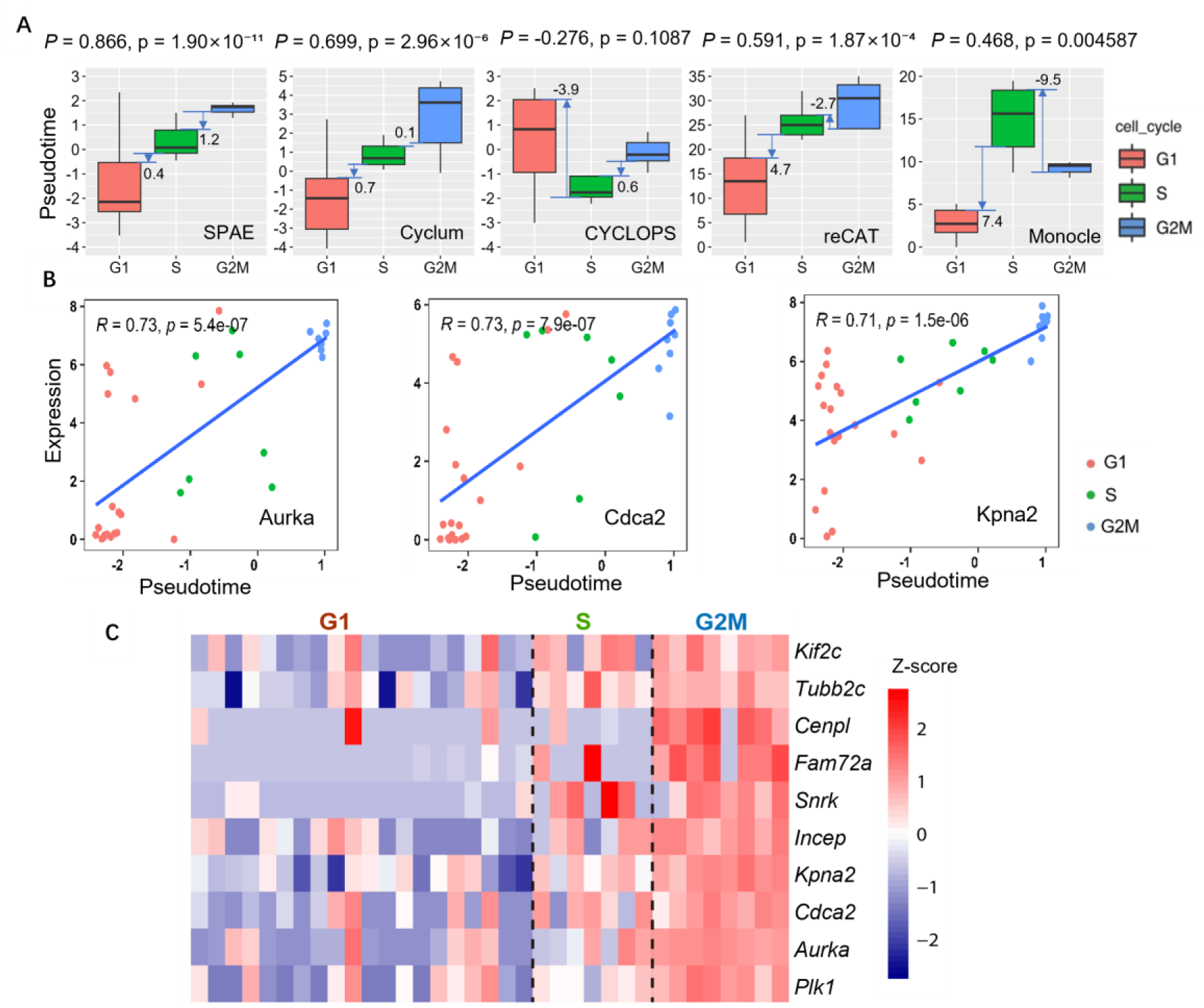
Cell cycle pseudotime analysis of mESCs Quartz-Seq data. **(A)** Boxplots show the distribution of cell cycle pseudotimes inferred by five different methods (SPAE, Cyclum, CYCLOPS, reCAT, and Monocle). The boxplots are colored according to the three stages of the cell cycle (G1, S, G2/M). **(B)** Correlation between the expression of three cell cycle marker genes (*Aurka*, *Cdca2*, *Kpna2*) and the cell cycle pseudotime estimated by SPAE. The top-left corner of each plot is marked with the correlation coefficient (R) and *P*-value, demonstrating a strong correlation between expression levels and pseudotime. **(C)** A heatmap displays several G2/M phase marker genes correlated with cell cycle pseudotime inferred by SPAE. Each column represents a cell. The color changes in the heatmap represent variations in gene expression levels, ranging from low (blue) to high (red).

### SPAE demonstrates superior accuracy and robustness in cell cycle characterization

To evaluate SPAE’s performance in predicting cell cycle progression, we follow the comparison strategy in Cyclum [20]. SPAE was compared with several models, including CCPE, Cyclum, CYCLOPS, cyclone, Seurat, and reCAT. The continuous pseudotime generated by SPAE, CCPE, Cyclum, and CYCLOPS was converted into discrete cell cycle phases using a three-component Gaussian mixture model. The performance was evaluated using seven classification metrics including Accuracy, Precision, Recall, F-score, Rand Index (RI), Normalized Mutual Information (NMI), and Adjusted Rand Index (ARI), across three datasets (mESCs Quartz-Seq, E-MTAB-2805 mESCs, and H1 hESCs). To account for the stochasticity of machine learning models, we evaluated each method ten times on each dataset and then calculated the average values for each performance metric. The radar plots demonstrate SPAE’s outstanding performance in the analysis of H1 hESCs dataset, with the clustering metrics achieving the highest values among all methods (**Figure 3A**, where 0.2, 0.4, 0.6, and 0.8 represent different thresholds). In the E-MTAB-2805 dataset analysis, SPAE led in all individual metrics, showcasing its superior performance compared to other models **(Figure 3B)**. Furthermore, in the analysis of the mESCs Quartz Seq dataset, SPAE continued to demonstrate strong performance (**Supplementary Figure S5)**, confirming its robustness across different datasets.

**Figure 3.**
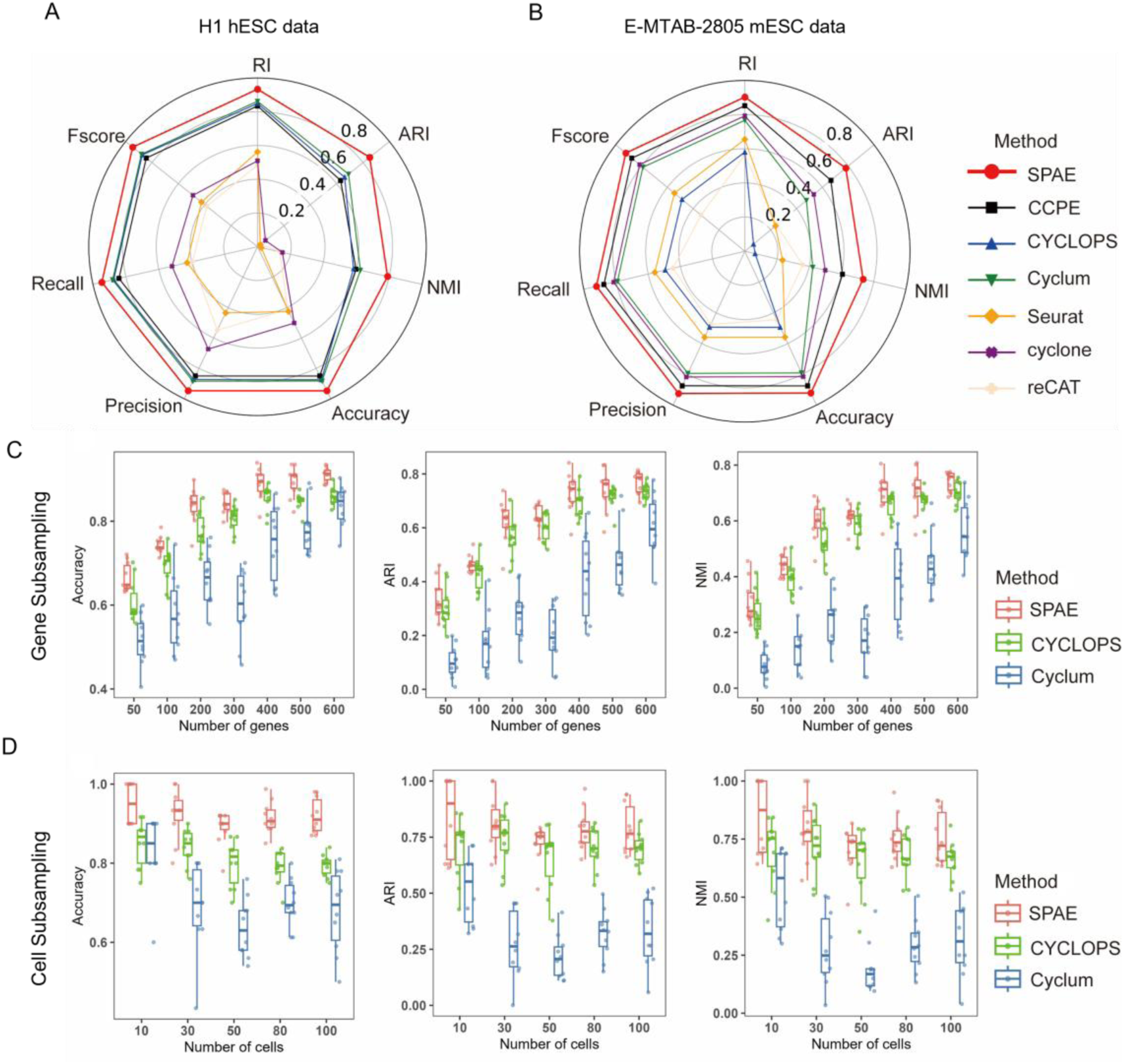
Inferring cell cycle stages from real datasets. **(A)** Radar chart showing seven multi-class classification metrics (Fscore, Recall, Precision, Accuracy, NMI, ARI, RI) used to evaluate the cell cycle classification accuracy of SPAE, CCPE, cyclone, Seurat, reCAT, Cyclum, and CYCLOPS on H1 hESCs data. **(B)** Radar chart displaying the same seven multi-class classification metrics for assessing the performance of SPAE and the six benchmark methods on E-MTAB-2805 mESCs data. **(C)** Robustness analysis under gene subsampling. The boxplots illustrate the Accuracy, Adjusted Rand Index (ARI), and Normalized Mutual Information (NMI) of SPAE, CYCLOPS, and Cyclum on datasets with varying numbers of genes (ranging from 50 to 600). **(D)** Robustness analysis under cell subsampling. The boxplots illustrate the performance of SPAE, CYCLOPS, and Cyclum in terms of Accuracy, ARI, and NMI metrics on datasets with varying cell counts (ranging from 10 to 100).

We further evaluated the performance of SPAE under different sample sizes by subsampling the scRNA-seq data from the H1 hESC dataset with fewer cells or genes. We conducted an analysis on seven sub-datasets with a diverse range of gene numbers, extending from 50 to 600, and five sub-datasets with a range of cellular numbers, extending from 10 to 100. Our results indicated a gradual increase in the median value of clustering metrics for SPAE, CYCLOPS and Cyclum with increasing number of genes (**Figure 3C, Supplementary Figure S6**). The findings of our analysis demonstrated the superiority of SPAE over CYCLOPS and Cyclum in terms of seven clustering metrics. In particular, SPAE exhibited a greater accuracy in predicting cell cycle stages using fewer genes compared to CYCLOPS and Cyclum. Moreover, SPAE displayed better performance with smaller numbers of cells compared to CYCLOPS and Cyclum. As the number of cells increased, the performance of SPAE declined gradually and reached a stable point (**Figure 3D, Supplementary Figure S7**). Conversely, the performance of Cyclum and CYCLOPS exhibited fluctuations but consistently remained below that of SPAE. However, we did not include cyclone, Seurat, and reCAT in Figures 3C-D due to their poor performance in the context of subsampling. When randomly sampling cells, the accuracy of all performance metrics for these methods was extremely low. Additionally, when randomly sampling genes, the number of selected genes was often too small, and many of the genes were not marker genes used by these models, resulting in their failure to produce any meaningful results. Therefore, we believe including these methods in the subsampling analysis would not provide valuable insights, as their performance was not comparable to SPAE, Cyclum and CYCLOPS under these conditions. Our analysis suggests that SPAE is more robust and exhibits higher prediction accuracy for sub-datasets with smaller numbers of genes or cells. A comprehensive comparison of the algorithmic features and architectures of these methods is provided in **Supplementary Table S1**.

### SPAE is robust to dropout events in the E-MTAB-2805 mESCs dataset

To assess SPAE’s robustness against dropout events in scRNA-seq, we utilized the E-MTAB-2805 mESCs dataset, which initially had a dropout rate of 24%. We first applied the MAGIC [24] model to impute the missing data, and subsequently introduced artificial dropout rates of 0%, 20%, 50%, and 70%. Our analysis indicated that SPAE’s performance was influenced by the dropout rate. However, the evaluation of the clustering metrics in **Figure 4** revealed that SPAE outperformed Cyclum, and CYCLOPS when the dropout rate was below 70%. At a dropout rate of 70%, all methods (SPAE, Cyclum and CYCLOPS) were no longer performant in estimating the cell cycle stages based on precision, recall and F1 score. For lower levels of dropout events, however, SPAE’s clustering metric values remained higher than those of Cyclum and CYCLOPS. Therefore, our findings suggest that the SPAE is in general more robust to dropout events compared to Cyclum and CYCLOPS, albeit at higher levels of dropout events none of the methods were able to obtain sufficient information to deliver performant cell cycle stage estimation.

**Figure 4.**
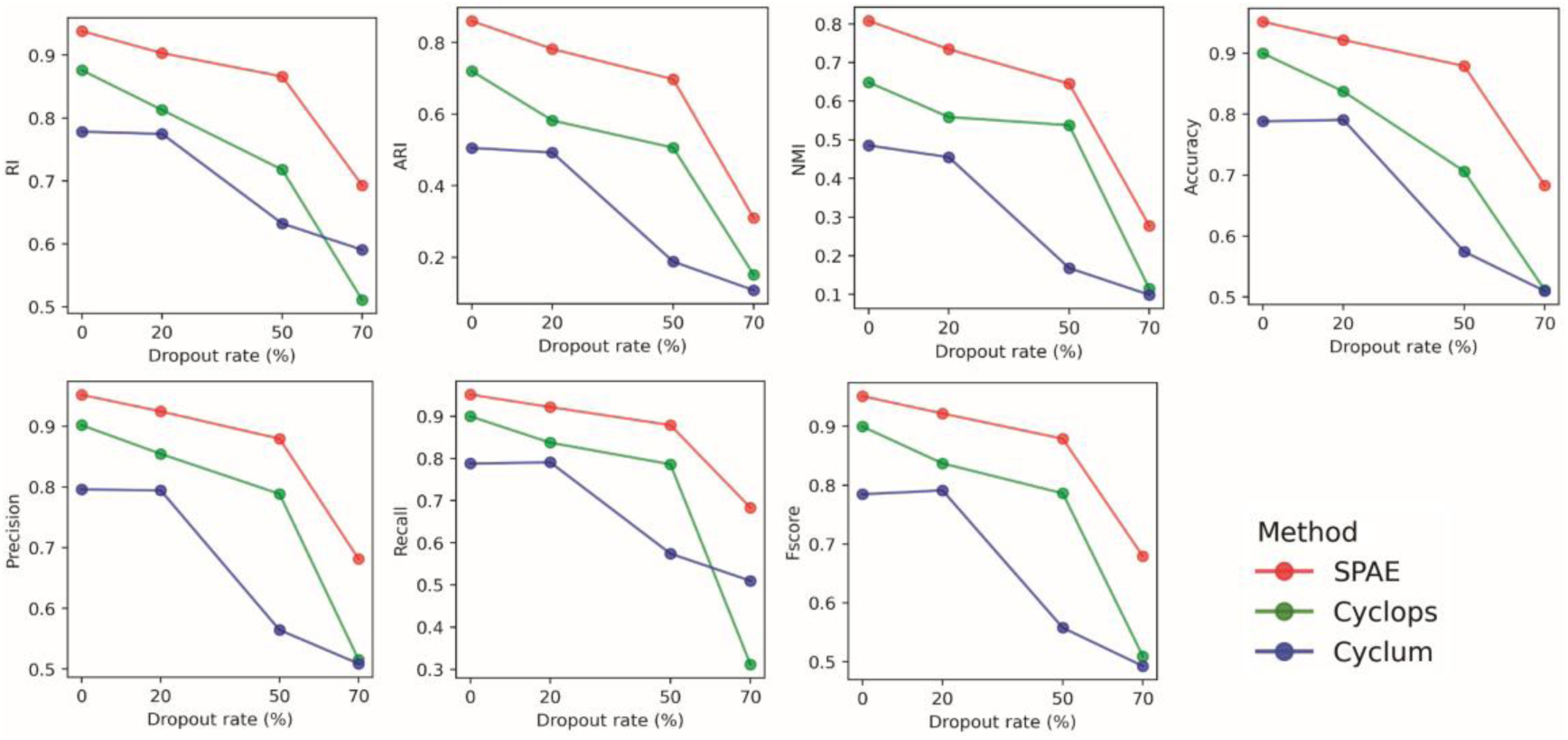
Robustness of SPAE in handling dropout events in the E-MTAB-2805 mESCs dataset. Evaluation of the impact of different dropout rates (ranging from 0% to 70%) on the performance of SPAE, Cyclum, and CYCLOPS. The comparison utilizes seven multi-class classification metrics: RI, ARI, NMI, Accuracy, Precision, Recall, and Fscore.

### SPAE identifies differentially expressed genes enriched in key cell cycle pathways

Performing differential gene expression analysis based on inferred cell cycle phases allows us to uncover variations in gene expression across distinct cell cycle stages. Using the DESeq2 package [34] within R/Bioconductor, we identified differentially expressed genes (DEGs) from cell cycle stages as inferred by SPAE. Subsequently, we compared these findings to those obtained from Cyclum, CYCLOPS, cyclone, Seurat, and reCAT using the E-MTAB-2805 mESCs dataset, applying stringent criteria (P.adjusted ≤ 0.05 and |log2FC| ≥ 1). Our gene set enrichment analysis[35] unveiled that DEGs identified by SPAE are predominantly related to cell cycle pathways and rank highly in enrichment analysis. Furthermore, these DEGs exhibited enrichment in biological processes closely associated with the cell cycle, encompassing pathways such as the p53 signaling pathway, progesterone-mediated oocyte maturation. Contrastingly, DEGs identified by Cyclum showed minimal relevance to the cell cycle, highlighting a significant difference (**Figure 5A**). CYCLOPS, cyclone, Seurat, and reCAT also exhibit some degree of cell cycle relevance, these methods are less effective and exhibit lower correlations compared to SPAE, failing to capture key pathways closely related to the cell cycle with the same level of significance (**Supplementary Figure S8**). We also explored the expression patterns of four genes enriched in the cell cycle pathway, *Cdc20*, *Fzr1*, *Cdk1* and *Ccnb1*, which are G2/M phase marker genes (**Figure 5B**). These genes were identified through a rigorous two-step selection process. First, differential gene expression (DEG) analysis was conducted using the DESeq2 package in R/Bioconductor, based on cell cycle stages inferred by SPAE from the E-MTAB-2805 mESCs dataset. Genes meeting the criteria of an adjusted P-value ≤ 0.05 and |log2FC| ≥ 1 were designated as DEGs. Second, gene set enrichment analysis of the DEGs revealed significant enrichment of *Cdc20*, *Fzr1*, *Cdk1*, and *Ccnb1* in both the cell cycle pathway and the progesterone-mediated oocyte maturation pathway. The critical role of these pathways in cell cycle regulation provided a strong rationale for their selection for further investigation.

**Figure 5.**
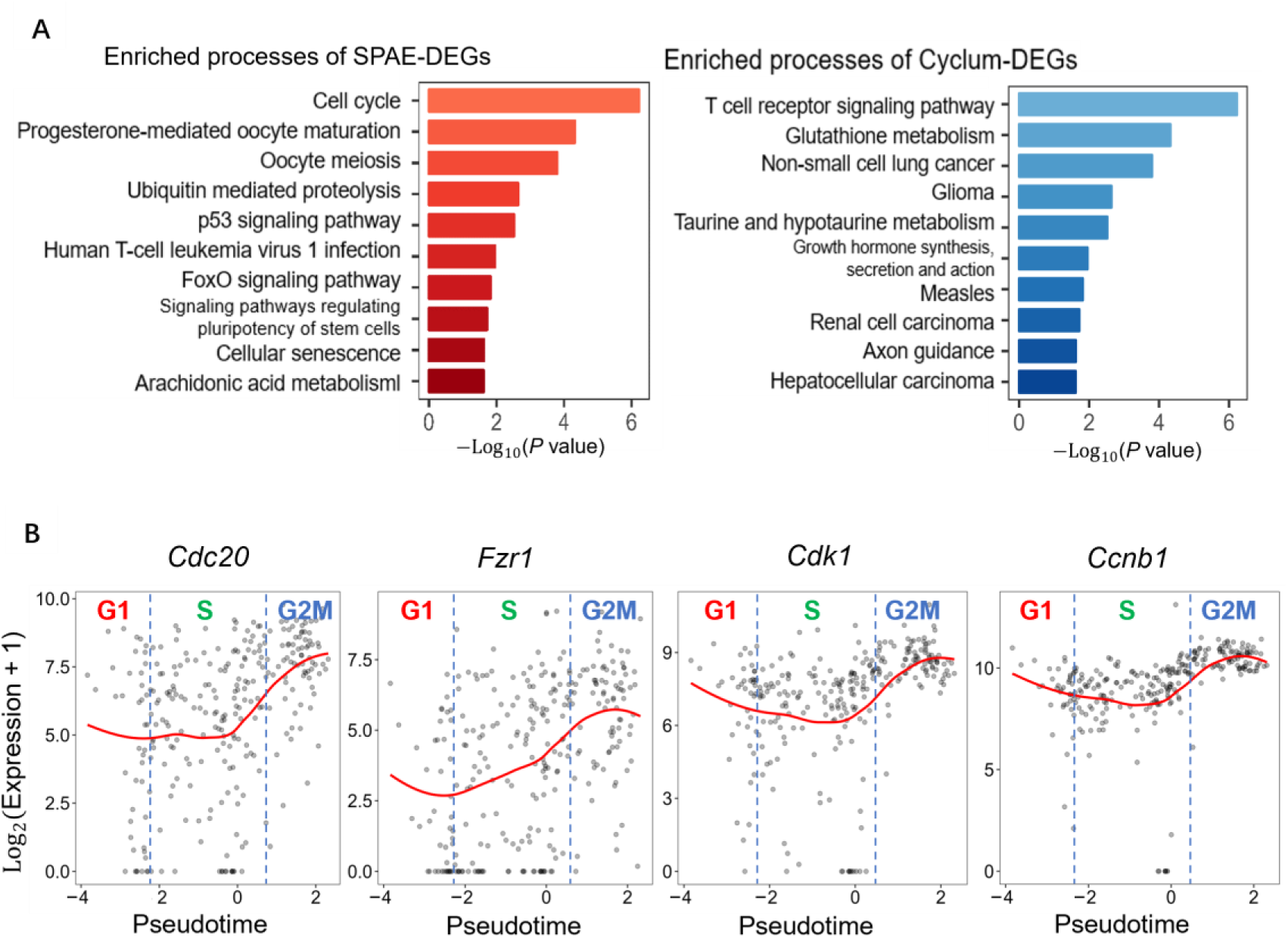
Differential gene expression analysis of E-MTAB-2805 mESCs Data. **(A)** Comparison of biological pathway enrichment. The left panel (red) shows the top ten biological processes enriched in DEGs identified by SPAE-inferred stages, including relevant terms like cell cycle, oocyte meiosis, and the p53 signaling pathway. The right panel (blue) shows processes associated with DEGs identified by Cyclum, which include unrelated pathways such as T-cell receptor signaling and non-small cell lung cancer. **(B)** Expression dynamics of G2/M phase marker genes. Scatter plots showing the expression variation of *Cdc20*, *Fzr1*, *Cdk1*, and *Ccnb1* along the cell cycle pseudotime inferred by SPAE. Each point represents a single cell, and the red curve indicates the smoothed gene expression trend along the inferred pseudotime.

### SPAE accurately detects Nutlin-induced G1 arrest in *TP53* wild-type cancer cells

To validate SPAE’s efficacy in cell cycle prediction, we analyzed a dataset of cancer cells treated with nutlin and DMSO. Nutlin, an antagonist of the MDM2-p53 pathway [36], is known to induce cell cycle arrest [37]. Nutlin promotes the stability and activity of p53 by inhibiting the interaction between MDM2 and p53, thereby inducing G1 phase arrest, particularly in *TP53* wild-type (WT) cells. As a key tumor suppressor protein, p53 initiates cell cycle checkpoints, especially the G1/S checkpoint, in response to cellular stress, preventing damaged cells from entering the S phase. Therefore, it is expected that Nutlin treatment in *TP53* WT cells will significantly increase the proportion of G1 phase cells and reduce the number of cells in the S and G2/M phases. We utilized two groups of data: one group consisted of cells treated with DMSO (the control group), and the other group comprised the same type of cells treated with nutlin [25]. The experimental samples comprised 7 *TP53* wild-type (WT). As shown in **Figure 6A**, SPAE’s predictions revealed a significant rise in G1 phase cells in *TP53* WT samples treated with nutlin, compared to the control group. Further analysis specifically targeting *TP53* WT cell lines reinforced this finding, showing a significant increase in the proportion of G1 phase cells, indicating that Nutlin induced G1 phase arrest in these cells (**Figure 6B**). Additionally, we identified DEGs for each cell cycle stage using DESeq2 (**Figure 6C**), revealing multiple pathways related to cell cycle regulation, particularly the regulation pathway of the G2/M checkpoint. In Nutlin-treated *TP53* WT cells, the G2/M checkpoint process ranks third among the DEGs. These findings not only confirm the high accuracy of SPAE in predicting cell cycle stages but also demonstrate its effectiveness in detecting G1 arrest induced by Nutlin in *TP53* WT cells.

**Figure 6.**
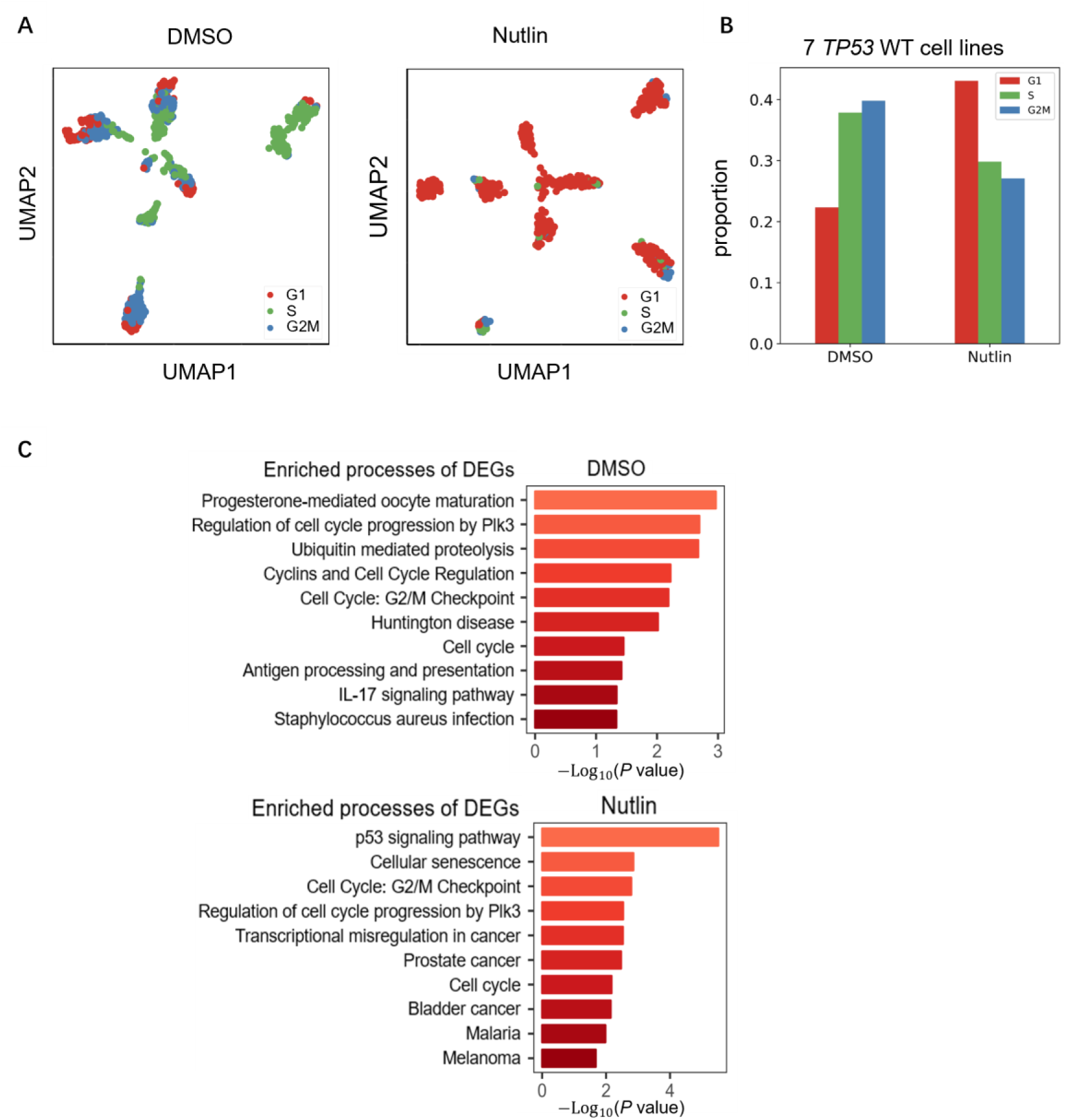
Validation of SPAE predictions using scRNA-seq datasets of cells treated with cell cycle perturbants. **(A)** UMAP visualization of 7 cancer cell lines treated with DMSO (control) or Nutlin. Cells are colored according to cell cycle phases (G1, S, G2/M) inferred by SPAE. **(B)** Quantitative analysis of G1 arrest. Bar chart showing the proportion of cells in different cell cycle stages for *TP53* wild-type (WT) cell lines in DMSO versus Nutlin treatment groups. SPAE accurately detects the accumulation of cells in the G1 phase upon Nutlin treatment. **(C)** Functional enrichment analysis. Bar charts showing the top enriched biological processes for Differentially Expressed Genes (DEGs) in the DMSO group (top) and the Nutlin group (bottom), highlighting pathways related to cell cycle regulation and p53 signaling.

### SPAE disentangles cell cycle effects from intrinsic cell states

In cellular biology, removing cell cycle effects is critical for accurately identifying cell types and understanding cellular differentiation [38]. Cells in different stages of the cell cycle often exhibit large differences in gene expression, which can obscure true biological signals and hinder functional analysis [39]. In this study, we evaluated the performance of SPAE in removing cell cycle effects across three datasets: mouse embryonic stem cells (mESCs), human myoblasts (hMyo), and breast cancer cells. The mESCs dataset includes data from four different withdrawal intervals of leukemia inhibitory factor (LIF) at 0, 2, 4, and 7 days. The hMyo dataset comprises scRNA-seq data from human myoblasts collected at various time points, namely 0, 24, 48, and 72 hours. The breast cancer dataset encompasses cell types with CSL gene knockout (CSLKO1 and CSLKO2) and wild-type controls (WT1 and WT2). To comprehensively assess SPAE’s performance in removing cell cycle effects, we compared it with four other methods: Cyclum, CCPE, ccRemover, and Seurat. We performed dimensionality reduction analyses using UMAP[40] on both raw data and the output of each method after cell cycle correction. The data were visualized with UMAP plots labeled by cell cycle stages (upper panels in **Figure 7A-C**) and by cell types, states or time points (lower panels in **Figure 7A-C**). The distribution of cells across different cell cycle stages was examined to evaluate the effectiveness of each method in mitigating cell cycle-driven clustering. Before the removal of the cell cycle effect, the distribution of cells in all three datasets was predominantly influenced by their cell cycle phases, obscuring the differences between cell types and hindering the clear separation of similar cell populations. However, after applying SPAE, cells of the same type, state or time point were more accurately clustered together, no longer being dispersed based on their cell cycle stages (**Figure 7A-C**). Specifically, in mouse embryonic stem cells, the raw data can distinguish cells from four different withdrawal intervals of leukemia inhibitory factor (LIF) at undifferentiated, 2, 4, and 7 days (**Figure 7A**). However, at each interval, cells in the three cell cycle stages are not well mixed, indicating that cell state classification is influenced by cell cycle effects. For instance, most ES7d cells are in the G1 phase. After applying SPAE to remove cell cycle effects, the distribution of cells across the three cell cycle stages in ES7d becomes more uniform. In contrast, CCPE effectively mixed cells from different cell cycle stages but failed to distinguish the four LIF withdrawal intervals. ccRemover distinguished the four intervals but did not evenly mix the three cell cycle stages, indicating residual cell cycle influence. Cyclum neither distinguished the LIF withdrawal intervals nor evenly mixed the three cell cycle stages. Seurat was able to distinguish between different LIF discontinuation intervals to some extent but failed to effectively and evenly mix cells in the three cell cycle stages (**Figure 7A**). Similar trends were observed in the hMyo dataset (**Figure 7B**) and the breast cancer dataset (**Figure 7C**). SPAE consistently outperformed other methods, effectively mixing cells across different cell cycle phases while maintaining distinctions between different time points, cell states, or cell types. This improved clustering suggests that SPAE successfully removed confounding cell cycle effects, allowing the data to reflect true biological differences. In contrast, other methods, including Cyclum, CCPE, Seurat, and ccRemover, did not achieve comparable results. Even after applying these methods, cell cycle-driven clustering remained evident (**Figure 7A-C**). SPAE ranked highest in both mixing cells from different cell cycle phases and maintaining clear separation of cells collected from different time points or biological states. These findings highlight SPAE’s robustness in removing cell cycle effects, making it a valuable tool for accurately analyzing single-cell transcriptomic data and uncovering true cellular identities.

**Figure 7.**
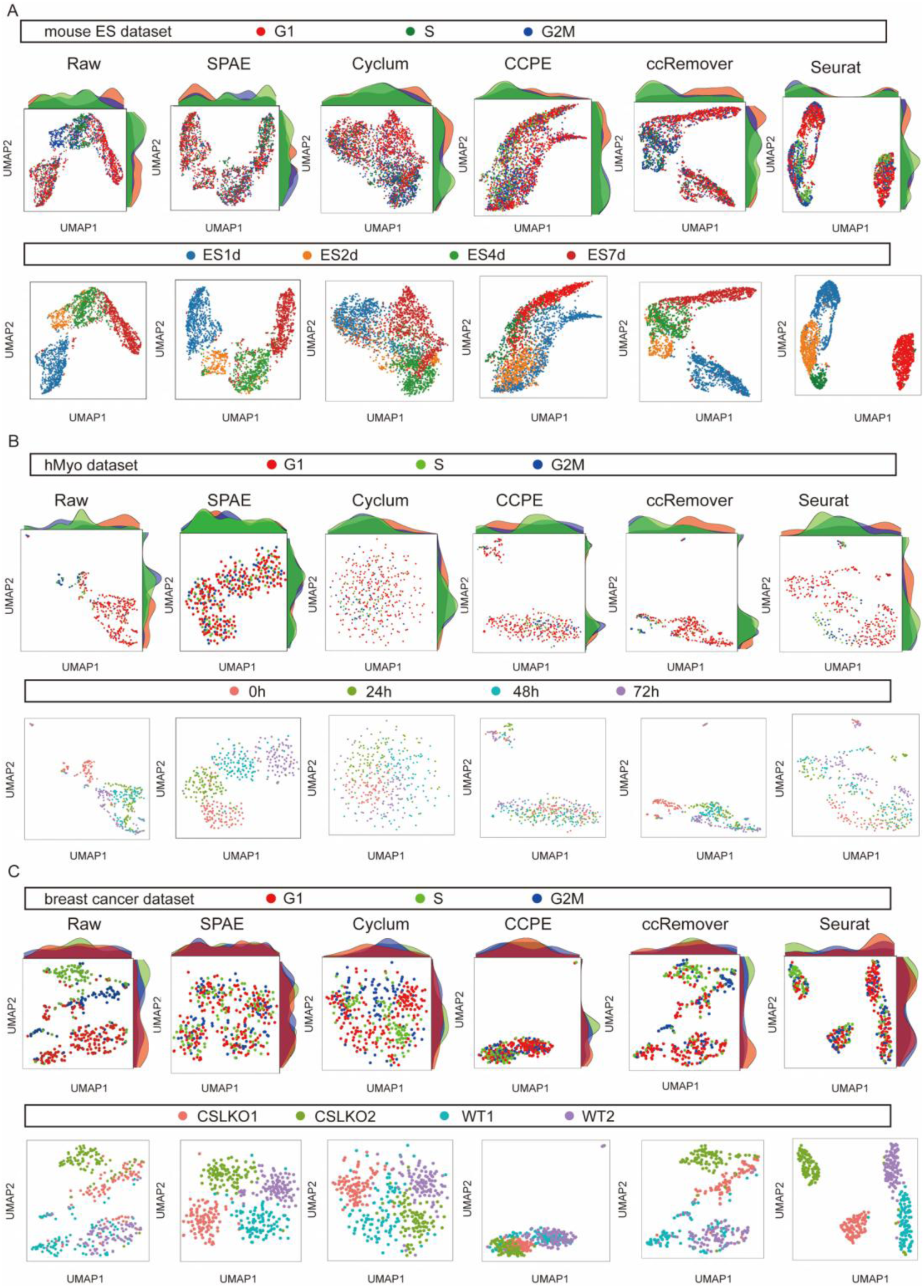
Comparison of different methods for removing cell cycle effects. **(A)** UMAP of mouse ES cells before and after removing cell cycle effects by SPAE, Cyclum, CCPE, ccRemover, Seurat. **(B)** UMAP of human myoblast cells before and after removing cell cycle effects by SPAE, Cyclum, CCPE, ccRemover, Seurat. **(C)** UMAP of breast cancer cells before and after removing cell cycle effects by SPAE, Cyclum, CCPE, ccRemover, Seurat. Raw represents data before cell cycle effect removal. Upper panel is labeled by cell cycle stages and lower panel is labeled by cell types, states or time points.

### Prediction of cell cycle transitions in breast cancer treatment using SPAE

Cell cycle dysregulation manifests as changes in the distribution of cells across various stages and alterations in the expression of cell cycle regulatory genes[41]. To assess SPAE’s effectiveness and discern changes in single-cell data, we analyzed scRNA-seq tumor data from 176,644 cells. These were divided into three treatment groups: endocrine therapy alone (letrozole plus placebo), intermittent high-dose combination therapy (letrozole plus ribociclib [600 mg/day, 3 weeks on/off]), and continuous low-dose combination therapy (letrozole plus ribociclib [400 mg/day])[26]. Patients underwent six cycles of treatment, with biopsies collected at baseline (day 0), the start of treatment (day 14), and at the end of treatment (around day 180 at surgery). SPAE was employed to predict transitions in the cancer cell cycle. Additionally, a cyclic generalized additive model was used to describe the dynamics of gene expression at various cell cycle stages. The SPAE-inferred cell cycle stages were used to color cells (**Figure 8A**). Reconstructing the cell cycle based on scRNA-seq data (**Figure 8A**), SPAE recovered the expected cell cycle stages, including the G1 checkpoint transition, where cyclin D initially rises, followed by *CDK6* expression. Additionally, we observed a decrease in *RB1* expression (a key G1 checkpoint protein) and an increase in *E2F3*, a proliferation gene. Moreover, based on SPAE-inferred cell cycle staging, we calculated the proportion of mitotic (S/G2 phase) cancer cells in each patient’s biopsy (**Figure 8B**). During combination therapy, an increase in the proportion of mitotic and proliferating (S/G2 phase) cancer cells was observed in each patient. Persistent tumors exhibited an increased frequency of proliferating cells, especially those undergoing high-dose combination therapy. In contrast, patients receiving only endocrine therapy showed fewer proliferating cells. These results suggest that in surviving subclonal populations, the ribociclib-enhanced G1/S checkpoint can be effectively bypassed. Subsequently, applying the cyclic generalized additive model revealed fluctuations in *ESR1* and *FOS* gene expression throughout the cell cycle. By applying this approach to cells sampled at different time points and patients undergoing different treatments, we differentiated whether treatment altered expression at specific cell cycle stages, or if gene expression was independent of cell cycle dysregulation. *ESR1* showed consistent expression levels throughout the cell cycle (**Figure 8C**). However, a decline in *ESR1* expression over time, coupled with an increase in *FOS* expression, was observed across the entire cohort receiving combination therapy (**Figure 8C**). Additionally, during combination therapy, a reduction in CDK inhibitor 2A (*CDKN2A* encoding p14 and p16) and an increase in *CDK6* expression from G1 to S/G2 phase were observed (**Figure 8C**). In summary, SPAE accurately estimates cancer cell cycle transitions and can be used to explore the phenotypic evolution of cancer cells under endocrine therapy and *CDK4/6* inhibitor treatment, as well as the relationship of these phenotypic changes to genomic variations. This information helps to reveal resistance mechanisms in early estrogen receptor-positive breast cancer and identify potential therapeutic targets.

**Figure 8.**
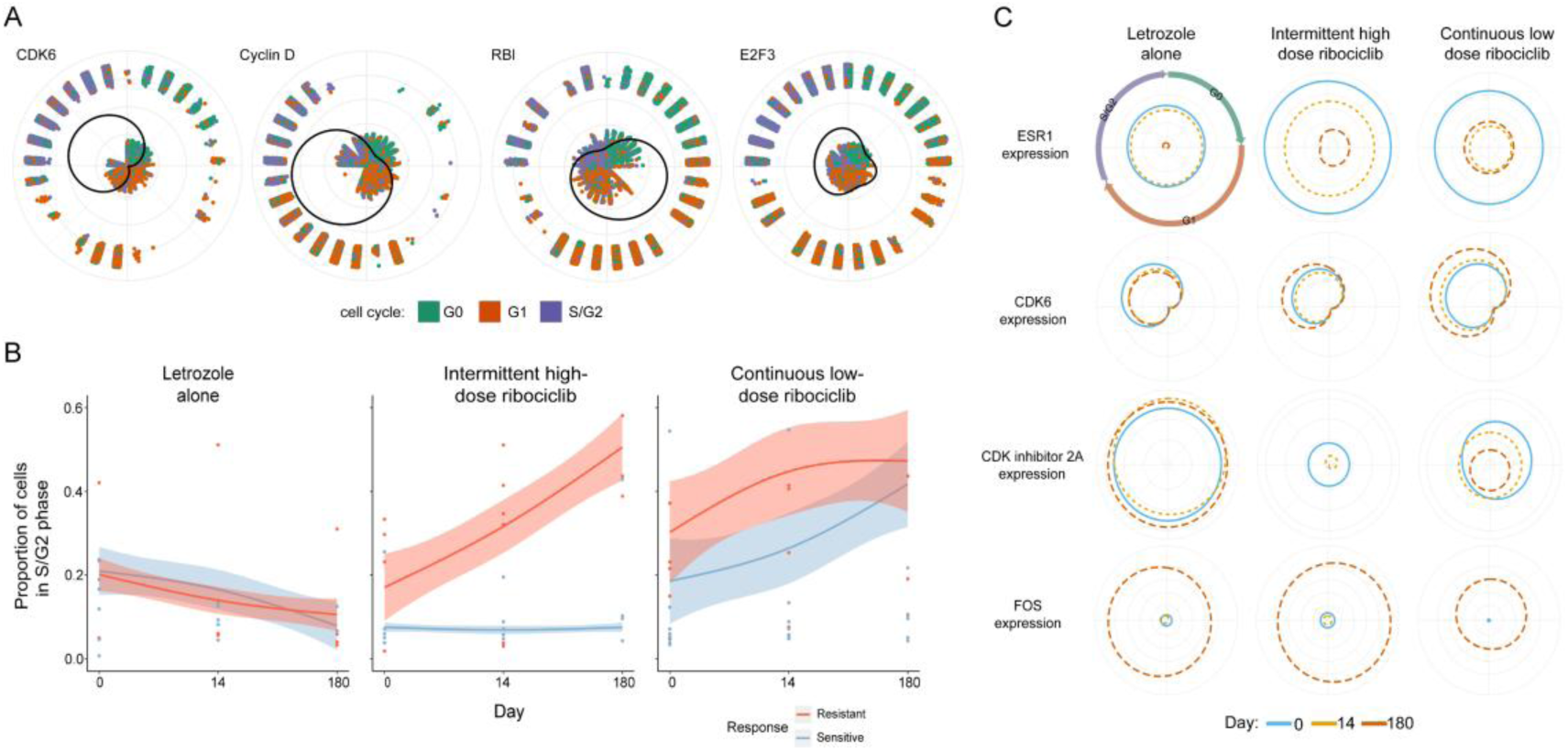
Analysis of cell cycle dysregulation based on SPAE. **(A)** Cell cycle phases colored based on predictions by SPAE, depicting expression changes of cell cycle regulatory genes *CDK6*, Cyclin D, *RB1*, and *E2F3* using a cyclic generalized additive model. The distance from the point to the origin indicates the expression of the cell at that phase. **(B)** Changes in the proportion of cells at different stages of the cell cycle (S/G2 phase) in patient tumor samples treated with Letrozole alone, intermittent high-dose Ribociclib, and continuous low-dose Ribociclib, as inferred by SPAE. Blue represents patients sensitive to treatment, and red represents patients resistant to treatment, with changes in cell proportions over time. **(C)** Changes in the expression of *ESR1*, *CDK6*, *CDKN2A*, and *FOS* before, during, and after the cell cycle and treatment (columns). Colored lines show the expected gene expression in cells throughout the cell cycle before (blue), during (orange), and after (red) treatment. The distance from the center of the circle indicates gene expression at a particular point in the cell cycle.

### SPAE identifies key transcription factors driving cell cycle transitions

The SPAE also enables us to identify potential transcription factors (TFs) responsible for the dynamics of gene expression along the cell cycle process. Transcription factors bind to specific DNA sequences (binding motifs) and activate the transcription of their target genes. They encode cellular programs for many functions required by the cell. We used SCENIC [42] to infer TF activity during the cell cycle. SCENIC is a method that computes gene regulatory networks in single-cell transcriptomic data through co-expression and motif analysis. Based on the cell cycle predicted by SPAE, we analyzed transcription factors using the mESCs Quartz-Seq dataset and the hESCs scRNA-seq dataset, with motif analysis predictions. In the mESCs Quartz-Seq dataset, observed TF activities suggested that the E2f family appears to be a group of key regulators, known to act at the onset of the cell cycle, especially during the G1/S transition [43, 44]. E2f1 and E2f2 peaked between the G1 and S phases, potentially activating genes required for the transition [45] (**Supplementary Figure S9A**). Specificity factor 1 (Sp1) and nuclear respiratory factor 1 (Nrf1) were both active in early G1 (**Supplementary Figure S9B**). For factors emerging from the hESCs scRNA-seq dataset, MYB is involved in the G2/M transition, functioning during the G2/M transition[46]. Kruppel-Like Factor 6 (KLF6), a transcription factor active in the G1 phase, can induce cell cycle arrest and reduce the rate of cell proliferation [47]. These results reveal the dynamic changes in transcription factor activity across different cell cycle stages, further supporting the critical role of transcription factors in regulating the cell cycle process. The observed TF activity patterns reflect the specific needs of cells at each stage for transcriptional regulation and offer a new perspective for understanding the cell cycle regulation of stem cells. These findings reinforce the central role of transcription factors in cell fate decisions and could provide potential targets for the treatment of cell cycle-related diseases.

## Discussion

In this work, we present SPAE, a computational framework that integrates a sinusoidal autoencoder with piecewise linear regression to decouple cell cycle dynamics from cell states. By utilizing an alternating optimization strategy, SPAE simultaneously learns continuous pseudotime and discrete cell clusters. While we employed a Gaussian Mixture Model (GMM) [48] to map pseudotime to discrete cell cycle phases (G1, S, G2/M), we acknowledge that standard GMMs may not fully capture the cyclic continuity and non-Gaussian patterns of biological processes. Nevertheless, this approach provides a practical approximation for delineating clinical stages from continuous trajectories.

Comprehensive benchmarking against established methods (including CCPE, Cyclum, CYCLOPS, Seurat and Monocle) across diverse scRNA-seq datasets demonstrated SPAE’s superior performance. Unlike linear models (e.g., CCPE) that struggle with complex nonlinear topologies, SPAE’s piecewise architecture effectively characterizes multi-stage transitions and nonlinear gene expression patterns. Consequently, SPAE exhibited greater accuracy and robustness in pseudotime inference, stability against data sparsity (dropout) and subsampling, and the identification of key cell cycle-related transcription factors. Critically, SPAE proved effective in removing cell cycle confounding effects, thereby revealing true cell type identities that were otherwise obscured. Biological validation on Nutlin-treated cancer cells further confirmed SPAE’s sensitivity in detecting specific G1 arrest and predicting therapy-resistant states, highlighting its translational potential.

Despite these advancements, limitations remain. Currently, SPAE lacks explicit modeling for complex technical variations, such as batch effects or tissue-specific biases, which may constrain its application in large-scale integrative studies. Future work will focus on incorporating mechanisms, such as adversarial domain adaptation, to mitigate these confounding factors and extend SPAE to complex disease models. In conclusion, SPAE provides a robust and flexible tool for dissecting the interplay between cell cycle dynamics and cell fate, offering new insights into biological heterogeneity and potential therapeutic targets.

## Code availability

SPAE developed for this study is implemented in Python 3.9 and is available for download on GitHub (https://github.com/YaJahn/SPAE). The code has also been submitted to BioCode at the National Genomics Data Center (NGDC), China National Center for Bioinformation (CNCB) (BioCode: BT008079), which is publicly accessible at https://ngdc.cncb.ac.cn/biocode/tool/ BT008079.

## CRediT author statement

**Jiahao Yi:** Methodology, Software, Formal analysis, Investigation, Writing – original draft. **Jiajia Liu:** Methodology, Software, Formal analysis. **Peng Guo:** Writing – original draft. **Yuan-Nong Ye:** Conceptualization, Supervision, Writing – review & editing. **Xiaobo Zhou:** Conceptualization, Supervision, Writing – review & editing. All authors read and approved the final manuscript.

## Competing interests

The authors declare no competing interests.

## Acknowledgements

This work was supported by the National Institutes of Health [R01LM014156, R01GM153822 and R01CA241930 to X.Z.], the National Science Foundation [2217515, 2326879 to X.Z.] and the National Natural Science Foundation of China [32160151 to Y.N.Y.].

## Supplementary material

Supplementary material is available at Genomics, Proteomics & Bioinformatics online.

## Supplementary Notes

**Supplementary Note 1: Piecewise linear regression model formulation.**

**Supplementary Note 2: Optimization of the autoencoder-based piecewise model.**

**Supplementary Table S1. Summary of computational methods for single-cell cell cycle analysis compared in this study.**

## Notes

https://github.com/YaJahn/SPAE

### Competing Interest Statement

The authors have declared no competing interest.

## References

1. Chang X, Zheng Y, Xu K. Single-Cell RNA Sequencing: Technological Progress and Biomedical Application in Cancer Research, Mol Biotechnol 2023; 66:1497–1519.

2. Jovic D, Liang X, Zeng H et al. Single-cell RNA sequencing technologies and applications: A brief overview, Clin Transl Med 2022;12:e694.

3. Wen L, Tang F. Recent advances in single-cell sequencing technologies, Precis Clin Med 2022;5:pbac002.

4. Papalexi E, Satija R. Single-cell RNA sequencing to explore immune cell heterogeneity, Nat Rev Immunol 2018;18:35–45.

5. Scialdone A, Natarajan KN, Saraiva LR et al. Computational assignment of cell-cycle stage from single-cell transcriptome data, Methods 2015;85:54–61.

6. Buettner F, Natarajan KN, Casale FP et al. Computational analysis of cell-to-cell heterogeneity in single-cell RNA-sequencing data reveals hidden subpopulations of cells, Nat Biotechnol 2015;33:155–160.

7. Trapnell C, Cacchiarelli D, Grimsby J et al. The dynamics and regulators of cell fate decisions are revealed by pseudotemporal ordering of single cells, Nat Biotechnol 2014;32:381–386.

8. Matthews HK, Bertoli C, de Bruin RAM. Cell cycle control in cancer, Nat Rev Mol Cell Biol 2022;23:74–88.

9. Bertoli C, Skotheim JM, de Bruin RA. Control of cell cycle transcription during G1 and S phases, Nat Rev Mol Cell Biol 2013;14:518–528.

10. Dang F, Nie L, Wei W. Ubiquitin signaling in cell cycle control and tumorigenesis, Cell Death Differ 2021;28:427–438.

11. Malumbres M, Barbacid M. Cell cycle, CDKs and cancer: a changing paradigm, Nat Rev Cancer 2009;9:153–166.

12. Mahdessian D, Cesnik AJ, Gnann C et al. Spatiotemporal dissection of the cell cycle with single-cell proteogenomics, Nature 2021;590:649–654.

13. Lee RD, Munro SA, Knutson TP et al. Single-cell analysis identifies dynamic gene expression networks that govern B cell development and transformation, Nat Commun 2021;12:6843.

14. Narotamo H, Fernandes MS, Moreira AM et al. A machine learning approach for single cell interphase cell cycle staging, Sci Rep 2021;11:19278.

15. Chen G, Ning B, Shi T. Single-Cell RNA-Seq Technologies and Related Computational Data Analysis, Front Genet 2019;10:317.

16. Liu J, Fan Z, Zhao W et al. Machine Intelligence in Single-Cell Data Analysis: Advances and New Challenges, Front Genet 2021;12:655536.

17. Butler A, Hoffman P, Smibert P et al. Integrating single-cell transcriptomic data across different conditions, technologies, and species, Nat Biotechnol 2018;36:411–420.

18. Liu Z, Lou H, Xie K et al. Reconstructing cell cycle pseudo time-series via single-cell transcriptome data, Nat Commun 2017;8:22.

19. Liu J, Yang M, Zhao W et al. CCPE: cell cycle pseudotime estimation for single cell RNA-seq data, Nucleic Acids Res 2022;50:704–716.

20. Liang S, Wang F, Han J et al. Latent periodic process inference from single-cell RNA-seq data, Nat Commun 2020;11:1441.

21. Anafi RC, Francey LJ, Hogenesch JB et al. CYCLOPS reveals human transcriptional rhythms in health and disease, Proc Natl Acad Sci U S A 2017;114:5312–5317.

22. Sasagawa Y, Nikaido I, Hayashi T et al. Quartz-Seq: a highly reproducible and sensitive single-cell RNA sequencing method, reveals non-genetic gene-expression heterogeneity, Genome Biol 2013;14:R31.

23. Leng N, Chu LF, Barry C et al. Oscope identifies oscillatory genes in unsynchronized single-cell RNA-seq experiments, Nat Methods 2015;12:947–950.

24. van Dijk D, Sharma R, Nainys J et al. Recovering Gene Interactions from Single-Cell Data Using Data Diffusion, Cell 2018;174:716–729.e727.

25. McFarland JM, Paolella BR, Warren A et al. Multiplexed single-cell transcriptional response profiling to define cancer vulnerabilities and therapeutic mechanism of action, Nat Commun 2020;11:4296.

26. Griffiths JI, Chen J, Cosgrove PA et al. Serial single-cell genomics reveals convergent subclonal evolution of resistance as early-stage breast cancer patients progress on endocrine plus CDK4/6 therapy, Nat Cancer 2021;2:658–671.

27. Klein AM, Mazutis L, Akartuna I et al. Droplet barcoding for single-cell transcriptomics applied to embryonic stem cells, Cell 2015;161:1187–1201.

28. Braune EB, Tsoi YL, Phoon YP et al. Loss of CSL Unlocks a Hypoxic Response and Enhanced Tumor Growth Potential in Breast Cancer Cells, Stem Cell Reports 2016;6:643–651.

29. McZgee VE, Carleton WT. Piecewise Regression, Journal of the American Statistical Association 1970;65:1109–1124.

30. Spearman C. The proof and measurement of association between two things. By C. Spearman, 1904, Am J Psychol 1987;100:441–471.

31. Roy A, Veroli MV, Prasad S et al. Protein Kinase D2 Modulates Cell Cycle By Stabilizing Aurora A Kinase at Centrosomes, Mol Cancer Res 2018;16:1785–1797.

32. Wang S, Cao K, Liao Y et al. CDCA2 protects against oxidative stress by promoting BRCA1-NRF2 signaling in hepatocellular carcinoma, Oncogene 2021;40:4368–4383.

33. Guo X, Wang Z, Zhang J et al. Upregulated KPNA2 promotes hepatocellular carcinoma progression and indicates prognostic significance across human cancer types, Acta Biochim Biophys Sin (Shanghai) 2019;51:285–292.

34. Love MI, Huber W, Anders S. Moderated estimation of fold change and dispersion for RNA-seq data with DESeq2, Genome Biol 2014;15:550.

35. Kuleshov MV, Jones MR, Rouillard AD et al. Enrichr: a comprehensive gene set enrichment analysis web server 2016 update, Nucleic Acids Res 2016;44:W90–97.

36. Shangary S, Wang S. Small-molecule inhibitors of the MDM2-p53 protein-protein interaction to reactivate p53 function: a novel approach for cancer therapy, Annu Rev Pharmacol Toxicol 2009;49:223–241.

37. Arya AK, El-Fert A, Devling T et al. Nutlin-3, the small-molecule inhibitor of MDM2, promotes senescence and radiosensitises laryngeal carcinoma cells harbouring wild-type p53, Br J Cancer 2010;103:186–195.

38. Barron M, Li J. Identifying and removing the cell-cycle effect from single-cell RNA-Sequencing data, Sci Rep 2016;6:33892.

39. Liu L, Michowski W, Kolodziejczyk A et al. The cell cycle in stem cell proliferation, pluripotency and differentiation, Nat Cell Biol 2019;21:1060–1067.

40. McInnes L, Healy J, Melville JJapa. Umap: Uniform manifold approximation and projection for dimension reduction 2018.

41. Pramparo T, Lombardo MV, Campbell K et al. Cell cycle networks link gene expression dysregulation, mutation, and brain maldevelopment in autistic toddlers, Mol Syst Biol 2015;11:841.

42. Aibar S, Gonzalez-Blas CB, Moerman T et al. SCENIC: single-cell regulatory network inference and clustering, Nat Methods 2017;14:1083–1086.

43. Tsai SY, Opavsky R, Sharma N et al. Mouse development with a single E2F activator, Nature 2008;454:1137–1141.

44. Gaubatz S, Lindeman GJ, Ishida S et al. E2F4 and E2F5 play an essential role in pocket protein-mediated G1 control, Mol Cell 2000;6:729–735.

45. Timmers C, Sharma N, Opavsky R et al. E2f1, E2f2, and E2f3 control E2F target expression and cellular proliferation via a p53-dependent negative feedback loop, Mol Cell Biol 2007;27:65–78.

46. Nakata Y, Shetzline S, Sakashita C et al. c-Myb contributes to G2/M cell cycle transition in human hematopoietic cells by direct regulation of cyclin B1 expression, Mol Cell Biol 2007;27:2048–2058.

47. Trucco LD, Andreoli V, Nunez NG et al. Kruppel-like factor 6 interferes with cellular transformation induced by the H-ras oncogene, FASEB J 2014;28:5262–5276.

48. Wan C, Chang W, Zhang Y et al. LTMG: a novel statistical modeling of transcriptional expression states in single-cell RNA-Seq data, Nucleic Acids Res 2019;47:e111.

