## Supplementary_Figures for "Deciphering Cell Cycle Dynamics and Cell States in Single-cell RNA-seq data with SPAE"

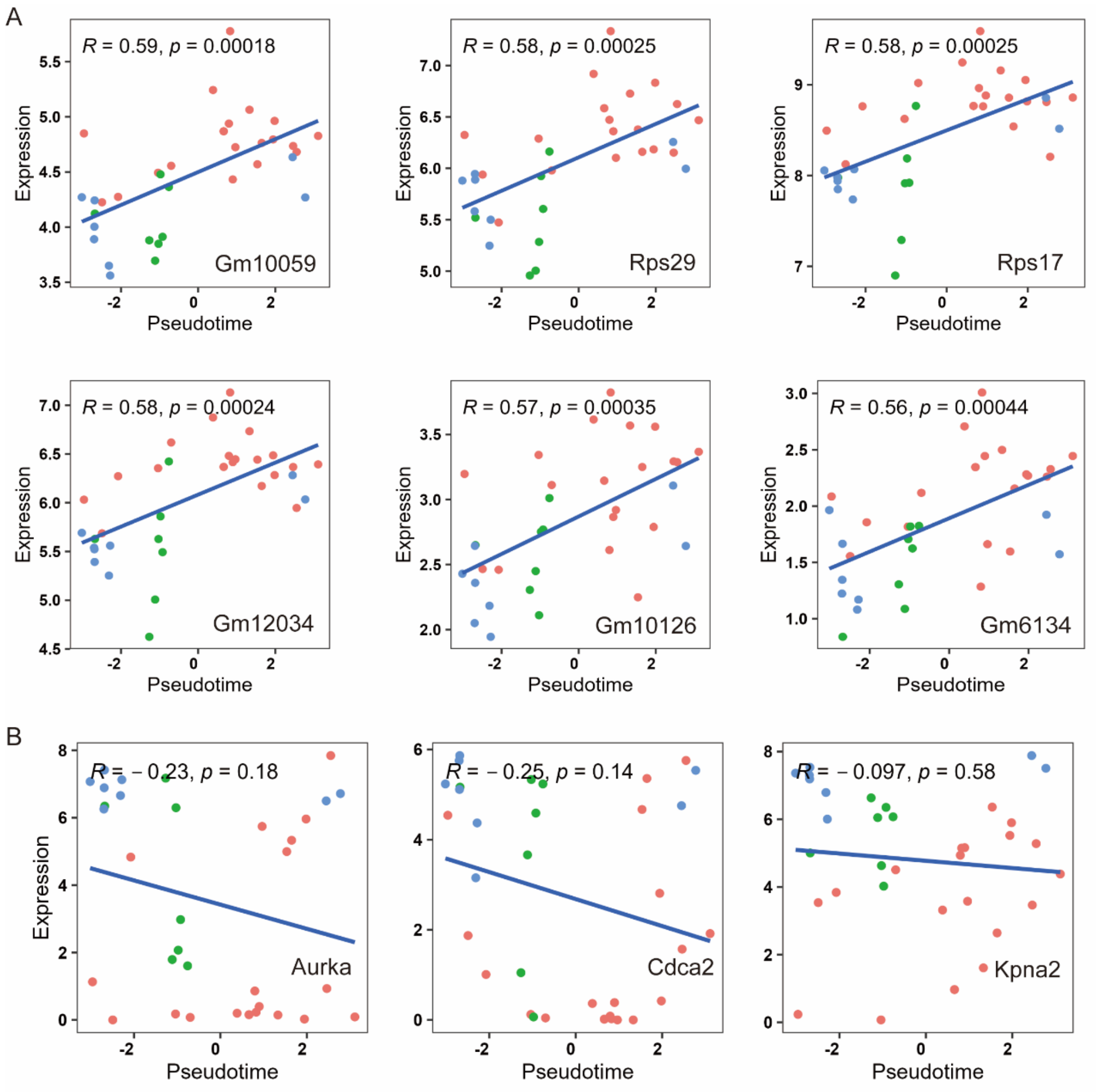


**Supplementary Figure S1**. **(A)** Pearson correlation between gene expression and pseudotime for the top six genes identified by CYCLOPS. **(B)** Pearson correlation between gene expression and pseudotime for *Aurka*, *Cdca2*, and *Kpna2* by CYCLOPS.

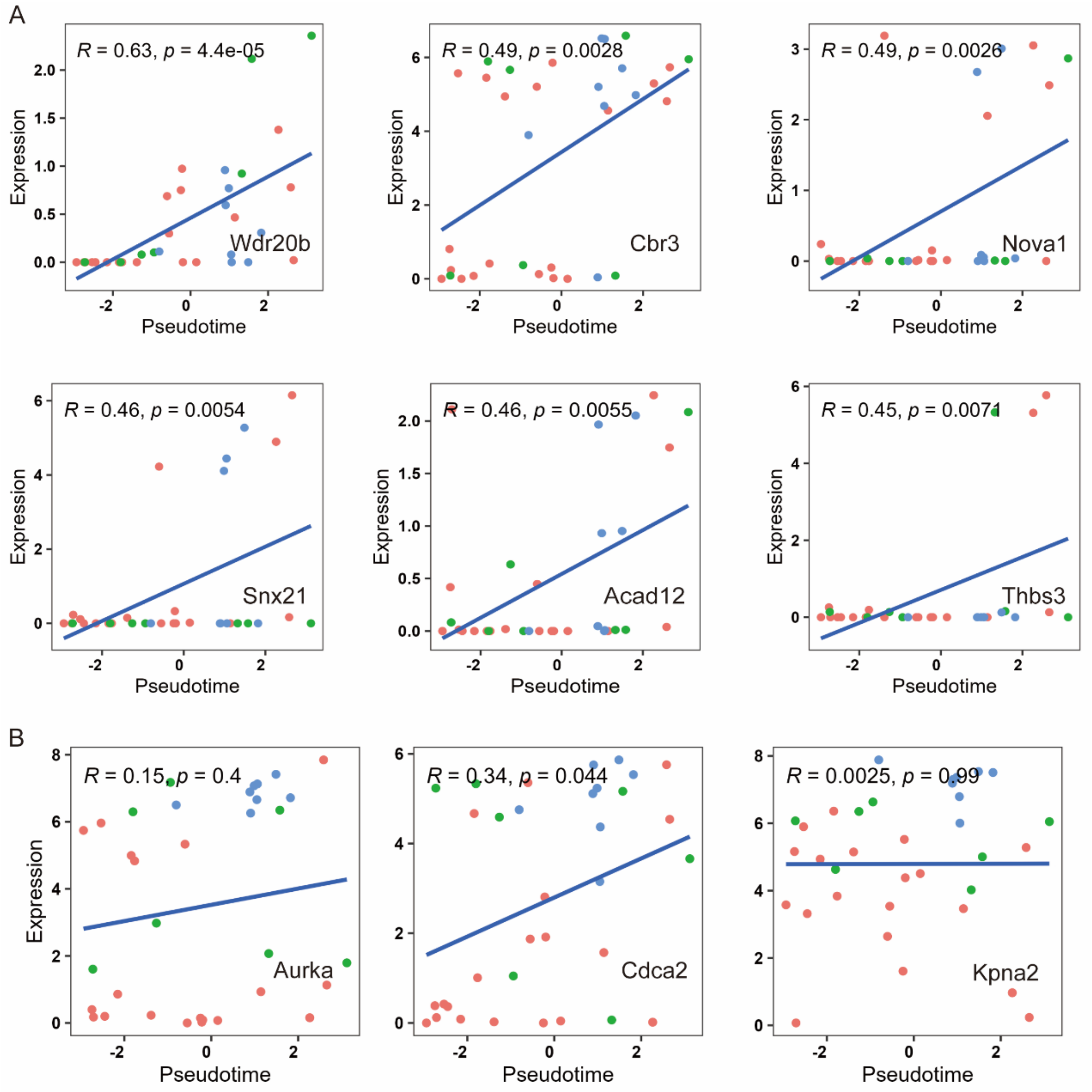


**Supplementary Figure S2**. **(A)** Pearson correlation between gene expression and pseudotime for the top six genes identified by Cyclum. **(B)** Pearson correlation between gene expression and pseudotime for *Aurka*, *Cdca2*, and *Kpna2* by Cyclum.

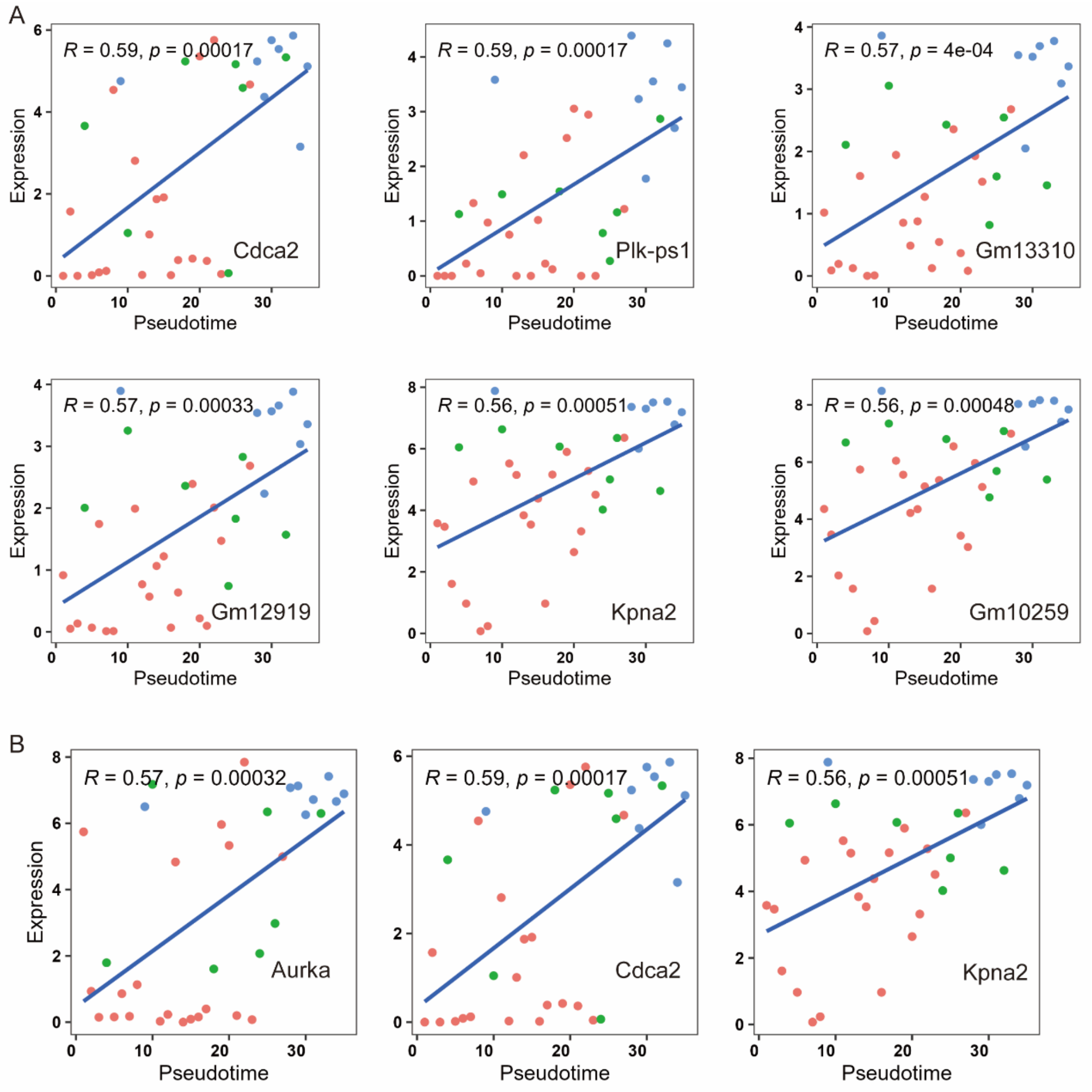


**Supplementary Figure S3**. **(A)** Pearson correlation between gene expression and pseudotime for the top six genes identified by reCAT. **(B)** Pearson correlation between gene expression and pseudotime for *Aurka*, *Cdca2*, and *Kpna2* by reCAT.

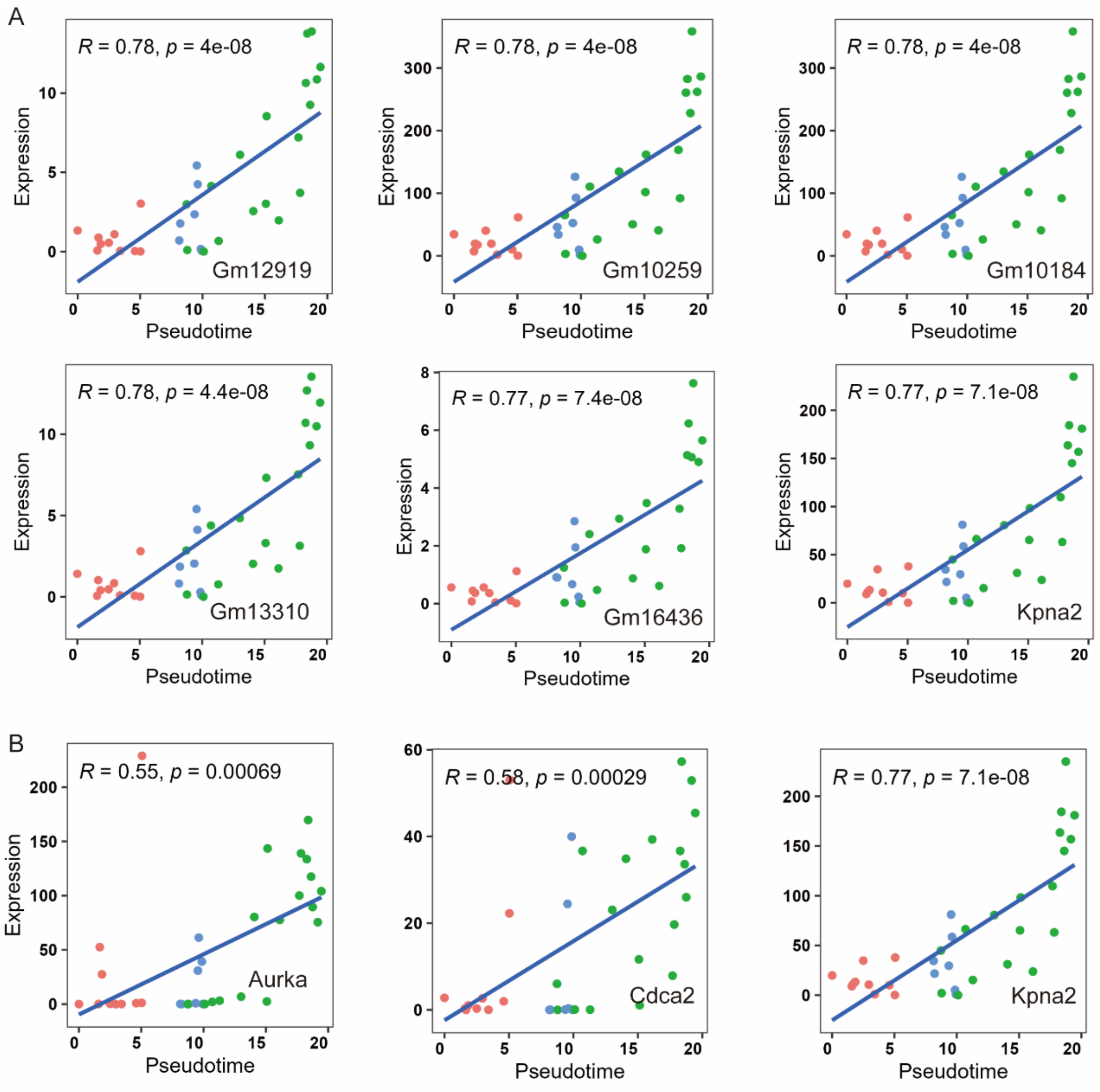


**Supplementary Figure S4**. **(A)** Pearson correlation between gene expression and pseudotime for the top six genes identified by Monocle. **(B)** Pearson correlation between gene expression and pseudotime for *Aurka*, *Cdca2*, and *Kpna2* by Monocle.

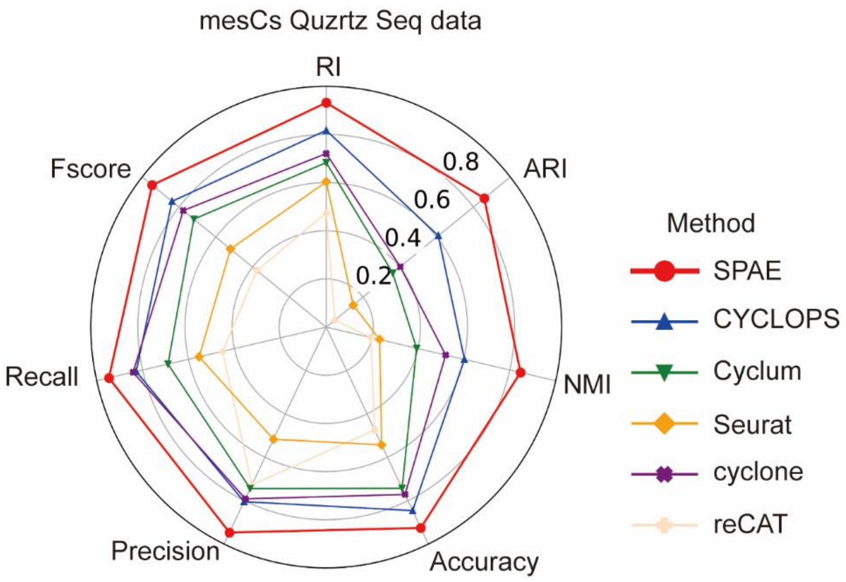


**Supplementary Figure S5**. Radar chart shows seven multi-class classification metrics used to evaluate the cell cycle classification accuracy of SPAE, cyclone, Seurat, reCAT, Cyclum, and CYCLOPS on mesCs Quartz Seq data.

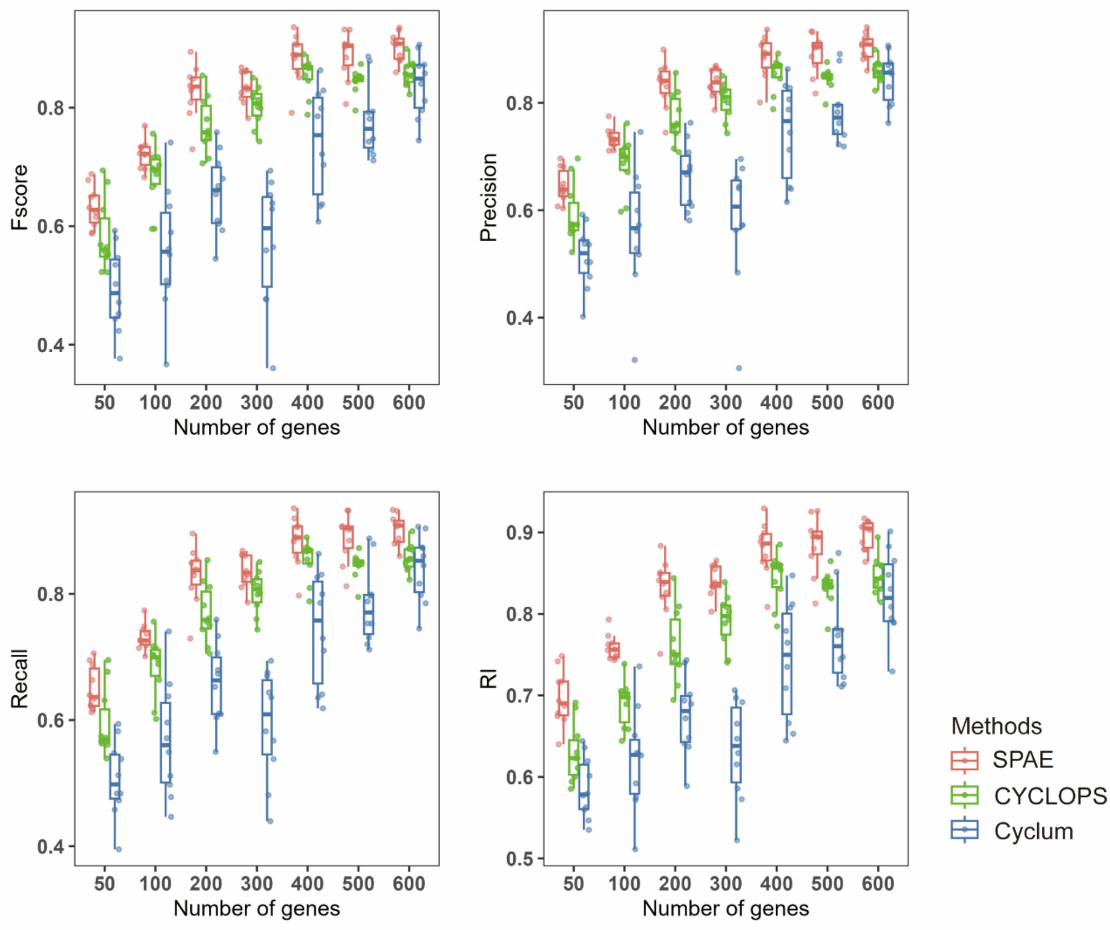


**Supplementary Figure S6**. Boxplots of Fscore, Precision, Recall and RI values indicate the performance of SPAE CYCLOPS and Cyclum on the subsampled datasets with different number of genes.

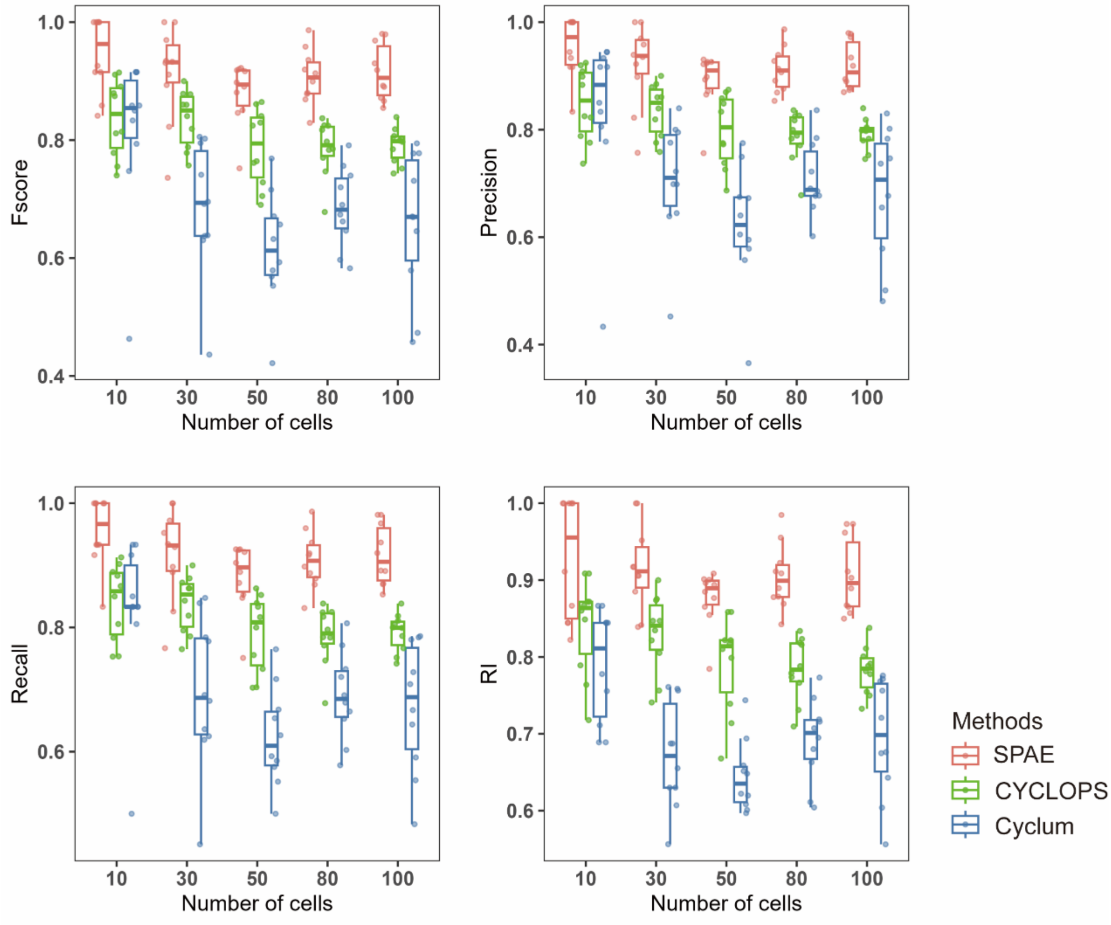


**Supplementary Figure S7**. Boxplots of Fscore, Precision, Recall and RI values indicate the performance of SPAE CYCLOPS and Cyclum on the subsampled datasets with different numbers of cells.

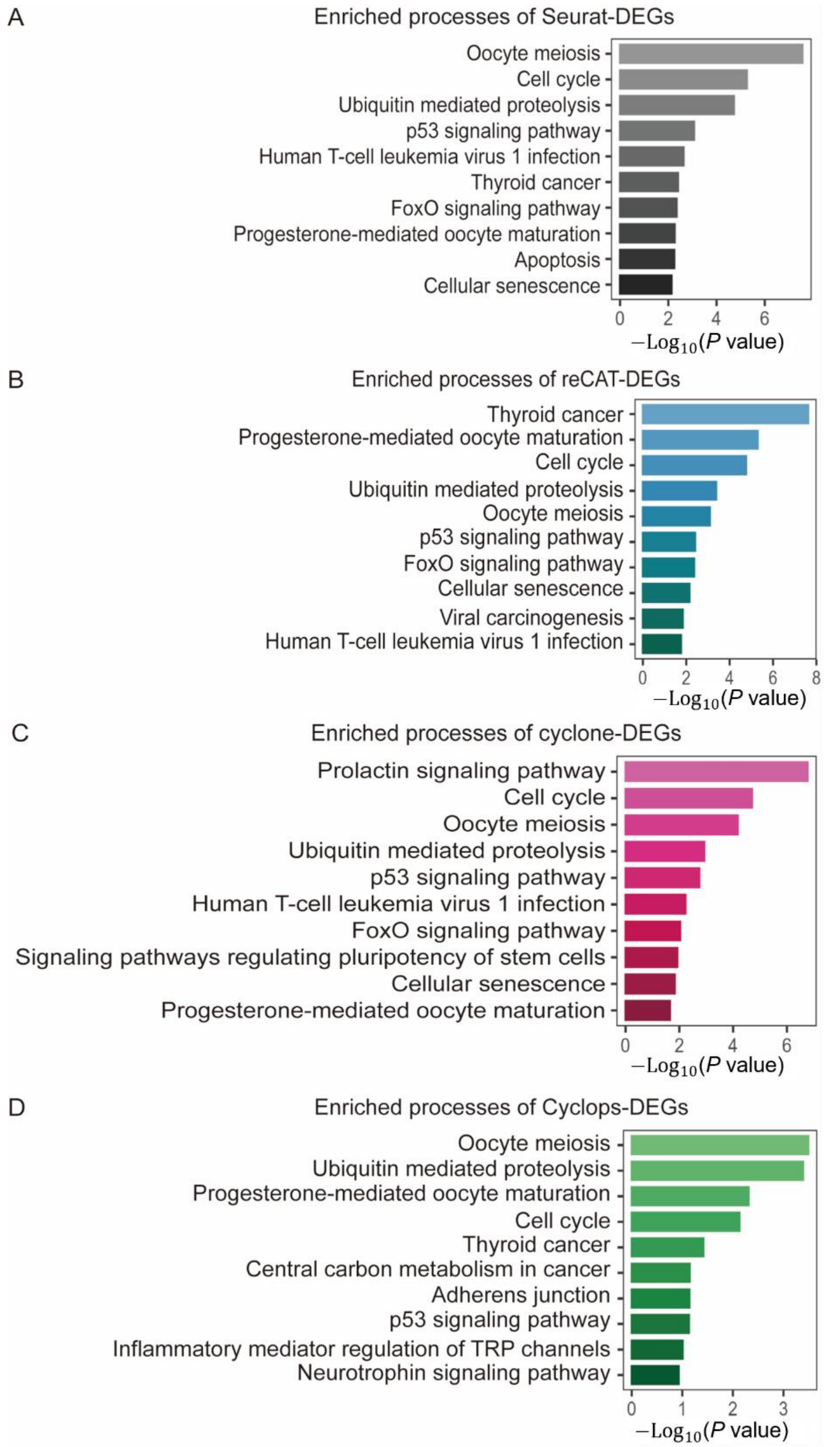


**Supplementary Figure S8**. Top ten enriched biological processes associated with DEGs identified by cell cycle stages inferred through Seurat, reCAT, cyclone and CYCLOPS.

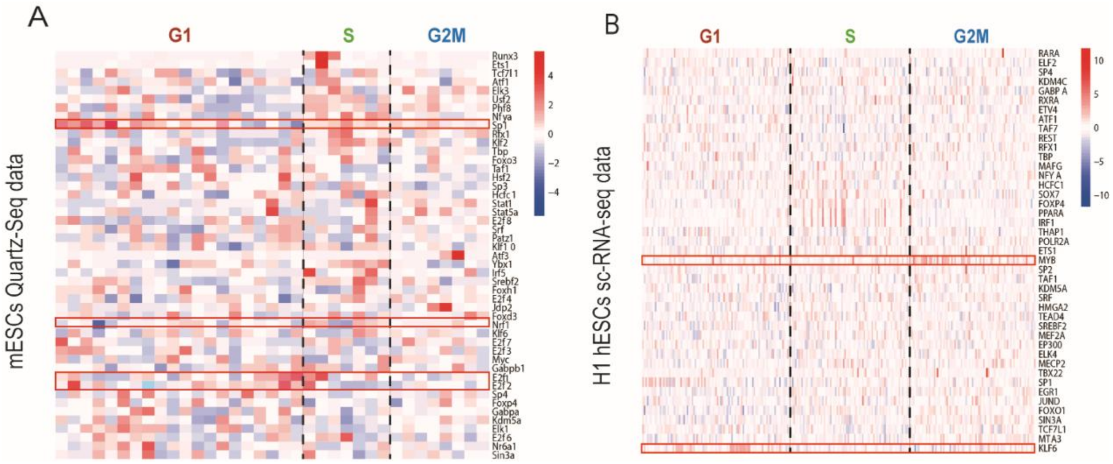


**Supplementary Figure S9.** SPAE predictions of core transcription factor activity during the cell cycle. **(A)** Motif activity of transcription factors in the cell cycle in the mESCs Quartz-Seq dataset. **(B)** Motif activity of transcription factors in the cell cycle in the hESCs scRNA-seq dataset. The color changes in the heatmap represent the variation in transcription factor activity, ranging from low (blue) to high (red).
