## Supplementary_Table for "Deciphering Cell Cycle Dynamics and Cell States in Single-cell RNA-seq data with SPAE"

**Supplementary Table S1. Summary of computational methods for single-cell cell cycle analysis compared in this study.**

| **Methods** | **Architecture** | **Focus** | **functions** |
|---|---|---|---|
| CCPE | 3D helical embedding with linear and nonlinear optimization | Captures continuous cyclic trajectories (linear assumption) | Reconstructs cell cycle progression from scRNA-seq data |
| cyclone | Pairwise gene ranking + SVM classification | Captures periodic structure based on known marker pairs | Assigns cell cycle phases (G1, S, G2M) |
| Seurat | Marker gene scoring (S.Score, G2M.Score) | Infers rhythmic transcriptomic variation via gene sets | Scores and classifies cell cycle phases |
| reCAT | Spectral clustering + Traveling Salesman Problem (TSP) | Recovers full cyclic ordering from unsynchronized data | Reconstructs temporal order of cycling cells |
| Cyclum | Autoencoder with sinusoidal activation (sin – cos latent space) | Infers nonlinear cyclic trajectories (Unsupervised) | Learns cyclic gene expression patterns in an unsupervised manner |
| CYCLOPS | Cyclic optimization with periodic basis decomposition | Orders cells along a closed elliptical curve | Estimates pseudotime along a cyclic trajectory |
| ccRemover | Principal component removal (unsupervised) | Focuses on eliminating cell cycle confounding effects | Removes cell cycle – associated variation from expression matrix |

Note: SVM, Support Vector Machine; TSP, Traveling Salesman Problem; scRNA-seq, single-cell RNA sequencing.
