## Supplementary_Text for "Deciphering Cell Cycle Dynamics and Cell States in Single-cell RNA-seq data with SPAE"

**Supplementary Note 1: Piecewise linear regression model formulation**

Piecewise linear regression is a regression method applied when the relationship between the dependent variable y and the independent variable x follows distinct linear trends over different ranges of x. This method uses indicator variables to fit the regression model for each segment (i.e., different ranges) of the data simultaneously.

In the context of our study, we model the cell state process as a piecewise linear relationship. Suppose there are $k$ cell types; the piecewise linear regression model is formulated as:

$$f(x) = f_0(x) + f_1(x)u(x - x_1) + f_2(x)u(x - x_2) + \cdots + f_k(x)u(x - x_k) \quad (1)$$

Where $f_k(x)$ represents the linear regression function for $k$th cell type and $u(x)$ is a gate function. The $k$th linear function is activated if the corresponding $u(x - x_k)$ equals to 1.

For example, considering a case with three distinct cell types, cells in each type are regressed along a linear function. The piecewise linear function is defined as:

$$\hat{y} = \begin{cases} w_1 x + b_1, & x < a_1 \\ w_2 x + b_2, & a_1 \le x < a_2 \\ w_3 x + b_3, & x \ge a_2 \end{cases} \quad \text{and} \quad \begin{cases} w_1 a_1 + b_1 = w_2 a_1 + b_2 \\ w_2 a_2 + b_2 = w_3 a_2 + b_3 \end{cases} \quad (2)$$

Where $w_1, w_2, w_3$ are the slope parameters for the linear functions corresponding to each cell type, dictating the rate of change in the dependent variable $y$. The intercept parameters $b_1, b_2, b_3$ represent the expected value of $y$ when $x$ equals zero for each respective piecewise of the model. The threshold parameters $a_1, a_2$ define the boundaries on the $x$ axis at which the slope and intercept parameters transition from one segment of the piecewise function to the next. $w_1, b_1$, $w_2, b_2$, $w_3, b_3, a_1, a_2$ are parameters to be optimized by solving the objective function

$$\min_{parameters} ||y - \hat{y}||^2 \quad (3)$$

**Supplementary Note 2: Optimization of the autoencoder-based piecewise model**

Optimizing the parameters of both the autoencoder and the piecewise linear model in SPAE presents a complex challenge. The autoencoder operates unsupervised, aiming to minimize reconstruction error, whereas the piecewise linear regression model typically requires supervised labels. To address this in our unsupervised framework, we employ pseudotime from the nonlinear component as a substitute label and utilize an alternating optimization strategy.

Pseudotime Initialization: Before the alternating process, we initialize the parameters of the nonlinear encoder ($w^{NL}$) using standard stochastic initialization. We then perform a forward pass to generate an initial estimation of pseudotime $z_n^c$. The pseudotime generated by this nonlinear part is used as the initial alternative label to start the piecewise component.

Alternating Optimization Strategy: The training process iterates through the following steps until convergence:

Step 1: Piecewise Regression and Threshold Optimization (Fix $z^c$, Optimize $w^p, a_i$):

Treating the current pseudotime $z^c$ as a fixed independent variable, we optimize the piecewise component. (1) Parameters: We optimize the weights ($w_i^p$) and biases ($b_i^p$) for each segment to minimize the regression error and the regularization term $\sum \alpha_i \parallel w_i^p \parallel^2$. (2) Thresholds ($a_i$): To optimize the transition thresholds $a_i$, we first order the cells based on their pseudotime $z^c$. We then systematically identify the optimal cut-points that minimize the sum of squared errors between adjacent linear segments, subject to the continuity constraints (Equation 2). Since the weight matrices are fixed during this step, the regularization terms become constants, reducing the optimization problem to minimizing the reconstruction loss.

Step 2: Autoencoder Refinement (Fix Piecewise, Optimize $w^{NL}, V$):

We fix the piecewise parameters ($w^p, b^p, a_i$) and update the nonlinear encoder ($w^{NL}$) and decoder ($V$) weights via backpropagation. This step minimizes the reconstruction loss along with the regularization terms ($\lambda \parallel w^{NL} \parallel^2 + \beta \parallel V \parallel^2$). Physically, this refines the mapping $x_n \rightarrow z_n^c$, adjusting the cell positions on the latent manifold to better align with the established piecewise linear structure.

This cycle repeats until the reconstruction error converges and the pseudotime ordering stabilizes.
